# The Intergenic Small Non-Coding RNA *ittA* is Required for Optimal Infectivity and Tissue Tropism in *Borrelia burgdorferi*

**DOI:** 10.1101/2020.02.24.962522

**Authors:** Diana N. Medina-Pérez, Beau Wager, Erin Troy, Lihui Gao, Steven J. Norris, Tao Lin, Linden Hu, Jenny A. Hyde, Meghan Lybecker, Jon T. Skare

## Abstract

Post-transcriptional regulation via small regulatory RNAs (sRNAs) has been implicated in diverse regulatory processes in bacteria, including virulence. One class of sRNAs, termed *trans*-acting sRNAs, can affect the stability and/or the translational efficiency of regulated transcripts. In this study, we utilized a collaborative approach that employed data from infection with the *Borrelia burgdorferi* Tn library, coupled with Tn-seq, together with borrelial sRNA and total RNA transcriptomes, to identify an intergenic *trans*-acting sRNA, which we designate here as *ittA* for *i*nfectivity-associated and *t*issue-*t*ropic sRNA locus *A*. The genetic inactivation of *ittA* resulted in a significant attenuation in infectivity, with decreased spirochetal load in ear, heart, skin and joint tissues. In addition, the *ittA* mutant did not disseminate to peripheral skin sites or heart tissue, suggesting a role for *ittA* in regulating a tissue-tropic response. RNA-Seq analysis determined that 19 transcripts were differentially expressed in the *ittA* mutant relative to its genetic parent, including *vraA, bba66*, *ospD* and *oms28* (*bba74*). Subsequent proteomic analyses also showed a significant decrease of OspD and Oms28 (BBA74) proteins. To our knowledge this is the first documented intergenic sRNA that alters the infectivity potential of *B. burgdorferi*.

**AUTHOR SUMMARY:** Lyme disease is a tick-borne infection mediated by the spirochetal bacterium, *Borrelia burgdorferi*, that is responsible for greater than 300,000 infections in the United States per year. As such, additional knowledge regarding how this pathogen modulates its regulatory armamentarium is needed to understand how *B. burgdorferi* establishes and maintains infection. The identification and characterization of small, non-coding RNA molecules in living systems, designated as sRNAs, has recalibrated how we view post-transcriptional regulation. Recently, over 1,000 sRNAs were identified in *B. burgdorferi*. Despite the identification of these sRNAs, we do not understand how they affect infectivity or *B. burgdorferi* pathogenesis related outcomes. Here, we characterize the *ittA B. burgdorferi* sRNA and show that it is essential for optimal infection using murine experimental infection as our readout. We also track the effect of this sRNA on the transcriptional and proteomic profile as the first step in providing mechanistic insight into how this important sRNA mediates its regulatory effect.

## INTRODUCTION

Lyme disease results from the infection by the spirochetal bacterium, *Borrelia burgdorferi,* and represents the most common vector-borne disease in the United States with an estimated 329,000 cases diagnosed each year [1]. *B. burgdorferi* is transmitted to mammalian hosts through the bite of infected *Ixodes* spp. ticks [2–4]. In humans, the infection is characterized by a flu-like illness and, in most instances, is accompanied by a skin lesion denoted as erythema migrans [5, 6]. The subsequent infection, if effectively treated with antibiotics early in the development of Lyme disease, can be cleared. If untreated, *B. burgdorferi* can disseminate throughout the host to distal organs and tissues resulting in multiple pathologies, including carditis, various neuropathies, and arthritis [5, 6].

*B. burgdorferi* oscillates in nature between vastly disparate environments of the tick vector and vertebrate hosts [2–4,7]. The enzootic life cycle of *B. burgdorferi* initiates by uninfected tick larvae feeding on an infected vertebrate (usually a small mammal or bird), resulting in the acquisition of the spirochete during the resulting blood meal [2–4]. The infected larvae then molt into a nymph and will seek another blood meal. It is at this point that *B. burgdorferi* infect vertebrate hosts, including dead end hosts such as humans. In order to survive in these changing host environments, *B. burgdorferi* alters its transcriptional and protein profiles [2,8,9]. Previous studies demonstrated that *B. burgdorferi* senses and responds to environmental cues such as temperature, pH, and dissolved gases, as well as unidentified host factors, to modulate its gene expression [10–19]. Despite significant insight into these processes, the mechanisms utilized by *B. burgdorferi* to modulate its gene expression for environmental adaptation continues to be an area of active research.

In *B. burgdorferi,* several transcriptional regulators have been identified and characterized, as well as a growing list of DNA interacting proteins, which serve to alter borrelial gene expression either directly or indirectly [3,20–33]. Many of these regulators govern, in part, the production of surface proteins involved in borrelial virulence [20,34– 37]. In addition to these regulators, recent results indicate that *B. burgdorferi* produces a battery of small non-coding RNA molecules, designated sRNAs [38–40]. The role and molecular mechanisms of the sRNAs in *B. burgdorferi,* or their impact on borrelial pathogenesis, is not well understood.

sRNA-mediated post-transcriptional regulation in bacteria commonly involves the alteration of transcript stability or translation efficiency; transcript targets include those that contribute to pathogenesis [41–43]. In addition, sRNAs can bind to proteins and modify protein activity [43–45]. As such, sRNAs encoded in intergenic regions of bacterial genomes are *trans*-acting regulators that can influence multiple, genetically unlinked transcripts [43,46–49]. These *trans*-acting sRNAs have partial complementarity to the transcripts they target. The resulting sRNA-mRNA duplex that forms can alter gene expression by multiple mechanisms including affecting mRNA stability, which can lead to the stabilization or the degradation of the mRNA [43,48,50–53]. The sRNA-mRNA duplex can also alter translation initiation, thus affecting translation efficiency [43,48,50–53]. The outcome of *trans*-acting sRNA gene regulation is an increase or decrease production of the encoded protein [43,48,50–53].

Recently, the sRNA transcriptome of *B. burgdorferi* was reported and 1,005 sRNA species were identified; many of these sRNAs are upregulated at 37°C, a condition that models the mammalian host temperature *in vitro* [40]. Independently, a Luminex-based procedure for the detection of Signature-tagged mutagenesis (STM) clones of the transposon (Tn) library of *B. burgdorferi* strain B31, identified several intergenic non-coding regions that, when genetically inactivated, exhibited infectivity deficits [54]. Here we further characterize one of these intergenic sRNAs, that maps to 17 kilobase linear plasmid (lp17) between *bbd18* and *bbd21,* which was designated as SR0736 [40]. We have renamed SR0736 as *ittA* for *i*nfectivity-associated and *t*issue-*t*ropic sRNA *A*. We independently inactivated this locus and showed that mutants lacking *ittA* are significantly attenuated and do not disseminate to additional skin sites or heart tissue. To our knowledge this is the first intergenic sRNA that has been linked to infectivity and pathogenesis of *B. burgdorferi*. Taken together, our data suggests that *ittA* exerts its effect by engaging several unlinked genetic targets and altering their production in a manner that is required for optimal infection and dissemination throughout the host.

## RESULTS

### Identification of sRNA associated with *B. burgdorferi* infectivity

Given the importance of sRNAs in other pathogenic bacteria [41, 43] and the population of sRNAs in *B. burgdorferi* that are induced at conditions that mimic the mammalian host temperature *in vitro* [40], we sought to determine if a subset of intergenic *trans*-acting sRNAs in *B. burgdorferi* contributed to borrelial pathogenesis. Initially, we utilized a *B. burgdorferi* transposon (Tn) mutant library [54], coupled with Tn-seq analysis following mouse infection [55, 56], to identify Tn insertions that mapped to intergenic (IG) regions or non-coding regions within the genome. We focused on intergenic Tn mutants that were represented in the initial *in vitro* grown inoculum used for infection but were substantially reduced following murine infection (Table 1). These data were overlapped with *B. burgdorferi* strain B31 sRNA annotations [40] to identify the Tn insertions that interrupted sRNAs. We identified eight such Tn mutants and tested their ability to establish infection as individual isolates by infecting C3H/HeN mice at a dose of 10^4^ *B. burgdorferi* cells. After 21 days a qualitative assessment of infectivity was determined (Table 2). Five of eight sRNA Tn mutants displayed reduced infectivity relative to the parent strain, 5A18NP1 [57], with a reduction in culture positive sites ranging from 25% to 75% (Table 2). After this initial screen, we focused on the sRNA (SR0736; [40]) that maps between genes *bbd18* and *bbd21* of linear plasmid 17 (lp17), which demonstrated the most significantly attenuated phenotype of the strains evaluated (Tables 1 and 2). Based on the phenotype observed for *B. burgdorferi* cells that have a Tn insertion in the SR0736 sRNA, we designated SR0736 as *ittA* for *i*nfection-associated and *t*issue-*t*ropic sRNA *A*.

**TABLE 1.** Reduced prevalence of candidate sRNA mutants during murine infection, based on Tn-seq^a^. Genetic mutations that either reduce (*guaA*, *dbpA*, *pncA*) or do not affect (*bbg22*, *bbk52*, *bbb28*) are included for comparison.

| Location of Tn insert | sRNA ID<sup>b</sup> | Input 1<sup>c</sup> | Input 2<sup>c</sup> | Input 3<sup>c</sup> | Output 1<sup>d</sup> | Output 2<sup>d</sup> | Output 3<sup>d</sup> | Avg.Output /Avg.Input | Comments |
| --- | --- | --- | --- | --- | --- | --- | --- | --- | --- |
| cp26; *guaA* (*bbb18*) | N/A<sup>e</sup> | 1.2032 | 0.7411 | 0.7736 | 0.0009 | 0.0018 | 0.0009 | 0.0013 | Control; attenuated |
| lp54; *dbpA* (*bba24*) | N/A<sup>e</sup> | 0.0041 | 0.0073 | 0.0084 | ND<sup>f</sup> | ND<sup>f</sup> | ND<sup>f</sup> | 0<sup>g</sup> | Control; attenuated |
| lp25; *pncA* (*bbe22*) | N/A<sup>e</sup> | 0.0089 | 0.0045 | 0.0027 | ND<sup>f</sup> | ND<sup>f</sup> | 0.0003 | 0.0185 | Control; attenuated |
| lp28-2; *bbg22* | N/A<sup>e</sup> | 0.0672 | 0.0991 | 0.0927 | 0.1688 | 0.2016 | 0.0267 | 1.53 | Unchanged |
| lp36; *bbk52* | N/A<sup>e</sup> | 0.0005<br>7 | 0.0002<br>8 | 0.0003<br>8 | 0.0012<br>6 | 0.0007 | 0.0005<br>1 | 2.01 | Unchanged |
| cp26; *bbb28* | N/A<sup>e</sup> | 1.5432 | 1.0517 | 1.1083 | 1.3479 | 3.5336 | 4.1401 | 2.44 | Unchanged |
| lp54; IG *bba34-bba36* | SR0897 | 0.0201 | 0.0210 | 0.0085 | ND<sup>f</sup> | ND<sup>f</sup> | ND<sup>f</sup> | 0<sup>g</sup> | 6.5x10<sup>5</sup> reads<sup>h</sup> |
| lp54; IG *bba66-bba68* | SR0912 | 0.1001 | 0.1327 | 0.2092 | 0.0000<br>5 | ND<sup>f</sup> | 0.0000<br>6 | 0.00025 | 6x10<sup>5</sup> reads<sup>h</sup> |
| cp26; IG *bbb03-bbb04* | SR0948 | 0.2269 | 0.1362 | 0.0982 | 0.0001 | ND<sup>f</sup> | 0.0006 | 0.0015 | 1.8x10<sup>4</sup> reads<sup>h</sup> |
| cp26; IG *bbb13-bbb14* | SR0962 | 0.0258 | 0.0107 | 0.0062 | ND<sup>f</sup> | ND<sup>f</sup> | ND<sup>f</sup> | 0<sup>g</sup> | 3x10<sup>6</sup> reads<sup>h</sup> |
| lp17; IG *bbd04-bbd05a* | SR0725 | 0.0378 | 0.0323 | 0.0145 | ND<sup>f</sup> | 0.0000<br>6 | ND<sup>f</sup> | 0.00071 | 2x10<sup>6</sup> reads<sup>h</sup> |
| lp17; IG *bbd18-bbd21* | SR0736/itt A | 0.0244 | 0.0549 | 0.0454 | 0.0001 | ND<sup>f</sup> | 0.0000<br>6 | 0.0013 | 1x10<sup>5</sup> reads<sup>h</sup> |
| lp28-3; IG *bbh36a-bbh36b* | SR0795 | 0.1628 | 0.2209 | 0.1344 | 0.0000<br>5 | ND<sup>f</sup> | 0.0000<br>6 | 0.00021 | 1x10<sup>5</sup> reads<sup>h</sup> |
| lp38; IG *bbj37-bbj41* | SR0869 | 0.0384 | 0.0456 | 0.0337 | ND<sup>f</sup> | ND<sup>f</sup> | 0.0000<br>6 | 0.00051 | 2.5 x10<sup>3</sup> reads<sup>h</sup> |

<sup>a</sup> Data following a 3 week infection with a library of Tn mutants [54]; <sup>b</sup> sRNAs from the *B. burgdorferi* sRNA library [40];
<sup>c</sup> % of Tn-seq reads from the libraries made from the inoculum used for mouse infection; <sup>d</sup> % of Tn-seq reads from the libraries made from bacteria recovered from infected mice; <sup>e</sup> Output is pooled data from 24 mice; <sup>f</sup> N/A = not applicable;
<sup>f</sup> ND = not detected; <sup>g</sup> indicates that no sequences were observed following Tn-seq; <sup>h</sup> represents rounded number of reads from sRNA-specific libraries [40].

Popitsch et al. identified *ittA* via deep sequencing and showed that the *ittA*-encoded sRNA was made at a higher level when *B. burgdorferi* is grown at 37°C [40]. The sRNA and 5’ end deep sequencing data indicate that *ittA* is a processed intergenic sRNA (Fig. 1) [40, 58]. The mapped primary 5’ ends are upstream from the *ittA* sRNA with minimal read coverage of the precursor RNA. In addition, the distal 5’ transcriptional start site for *ittA* overlaps with the transcription start site for *bbd18* on the opposite DNA strand (Fig. 1).

**Figure 1.**
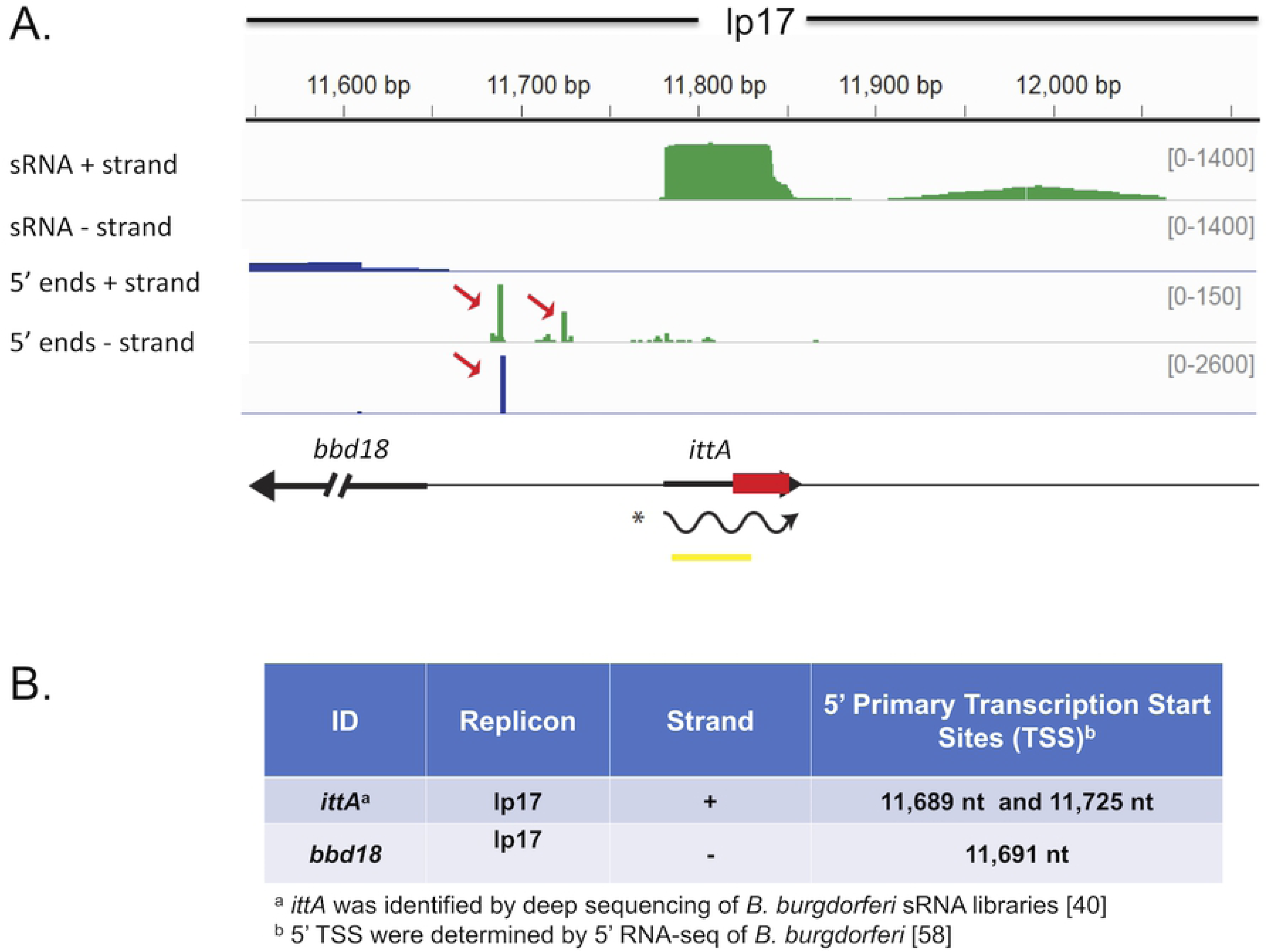
Location of SR0736/*ittA* sRNA on linear plasmid 17 (lp17). The deep sequencing results of the *ittA* sRNA are displayed in a coverage map. The negative strand coverage is shown in blue and the positive strand coverage in green. The genomic context is illustrated below the coverage maps: black arrows indicate the annotated ORFs, the yellow box indicates the Northern probe used, and the wavy line is the proposed mature sRNA species as determined by RNA-seq and Northern blot analysis. The red box represents the antibiotic resistance marker that is inserted into the *ittA* sRNA locus to obtain strain DM103. The sRNA is encoded in the positive strand and has two primary 5’ ends indicating transcriptional start sites (TSS) denoted by red arrows in the 5’ end + track. One of the TSS overlaps with the *bbd18* TSS, observed in the 5’end - track. The sequencing results show the sRNA processed into its mature form.

### Genetic inactivation of the *ittA* sRNA in *B. burgdorferi*

To independently test the role of the *ittA* sRNA in borrelial pathogenesis, we genetically interrupted *ittA* as depicted in Fig. 2A. The parent strain is the B31 derivative ML23 that lacks the 25 kb linear plasmid [59]. Due to the absence of lp25, this strain is non-infectious in the murine model of experimental infection. However, when a region of lp25, containing the *bbe22* gene, is provided in *trans* infectivity is restored and provides selective pressure for the maintenance of plasmids encoding it during experimental infection [60, 61]. The shuttle vector pBBE22*luc* used here contains *bbe22* and the firefly luciferase reporter that facilitates bioluminescent imaging as reported previously [60,62,63].

**Figure 2.**
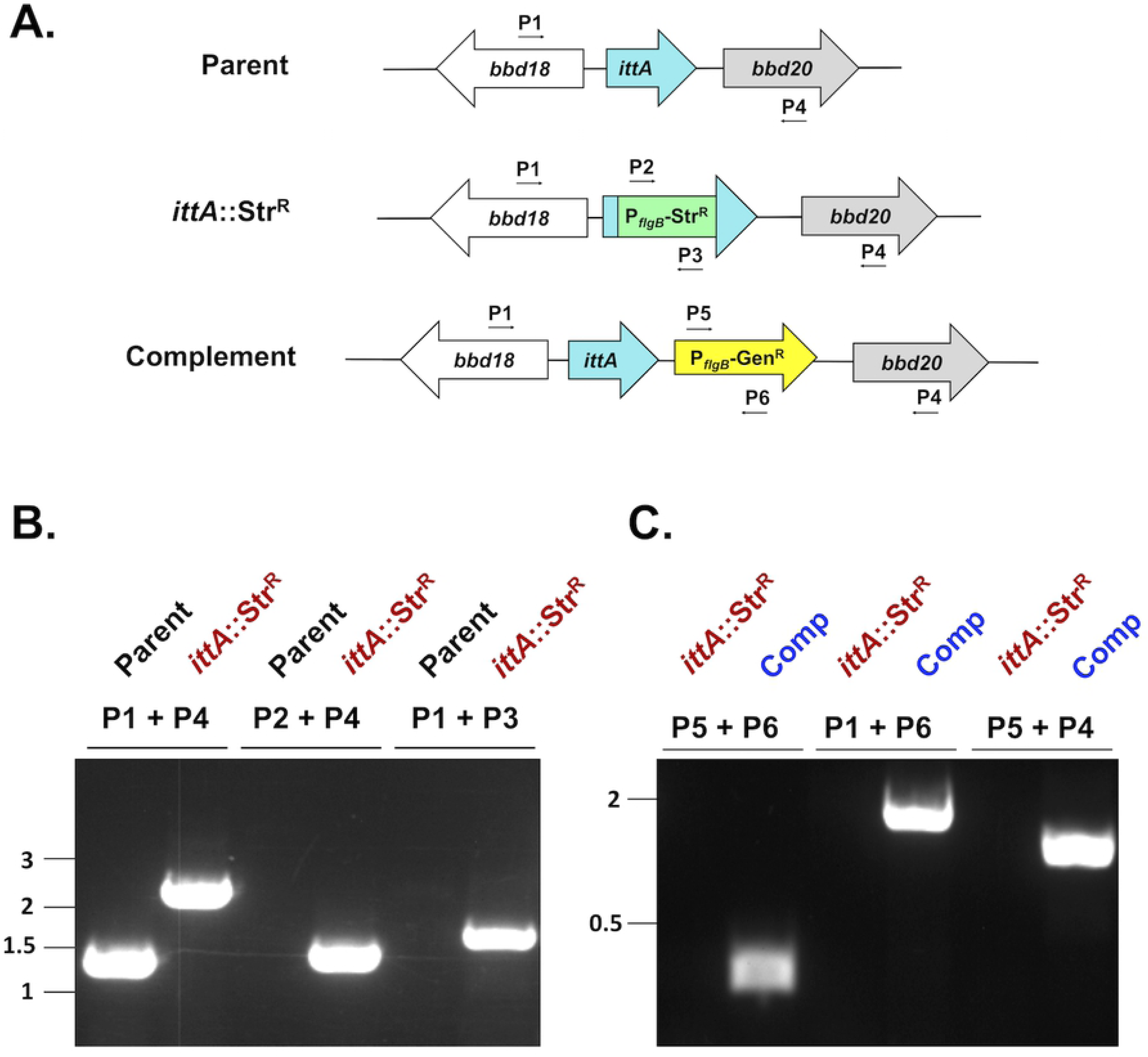
Strategy and confirmation of the insertional inactivation of the *ittA* sRNA. A. Schematic representation of *ittA* insertional inactivation strategy. The region of lp17 from the parent strain (ML23) is shown at the top. A P*_flgB_*_-_Str^R^ cassette was inserted into the 3’ end of the *ittA* sRNA and the mutant (DM103) was obtained following homologous recombination. Following isolation of the mutant strain, the *ittA* sRNA locus was reintroduced in the complemented strain (DM113). B. Primer pairs P1/P4, P2/P4 and P1/P3 (Table S7) were used to confirm the presence of the P*_flgB_*_-_Str^R^ in the mutant (*ittA*:Str^R^), relative to its parent strain by PCR. C. For the *cis* complementation of the *ittA* sRNA mutant strain, the P*_flgB-_*Str^R^ cassette and the flanking region were replaced by the *ittA* sRNA sequence linked to a P*_flgB_*_-_Gent^R^ cassette by a double crossover homologous recombination event. The resulting complement strain (Comp) was evaluated by PCR using primer pairs P5/P6, P1/P6 and P5/P4 (Table S7). Note that the distances between genetic loci are not shown to scale.

The *ittA*-encoding sRNA was insertionally inactivated in the *B. burgdorferi* strain ML23 (Fig. 2A). Transformants were selected with streptomycin and PCR was employed to distinguish the parent from potential *ittA*::Str^R^ mutant candidates (Fig. 2B). The resulting *ittA* mutant strain was designated DM103. We then genetically restored the *ittA* sRNA in *cis* in strain DM103 by selecting for resistance to gentamicin and then screening for sensitivity to streptomycin (Fig. 2A). Candidates predicted to encode *ittA* were vetted further using PCR (Fig. 2C). The *ittA* complement strain was designated DM113. Following the aforementioned screen of the *ittA*::Str^R^ mutant strain DM103 and the complement strain DM113, both were transformed with pBBE22*luc* so they could be tested in the murine experimental model of infection using a firefly luciferase reporter [60,62,63]. Note that all strains maintained plasmid content identical to the parent strain (data not shown).

To confirm that the *ittA* mutant and complement strains lacked and restored the *ittA* sRNA, respectively, both Northern blot and Reverse Transcriptase PCR (RT-PCR) were performed (Fig. 3). Notably, the *ittA* sRNA was detected in the parent strain ML23 and the complement strain DM113 (Comp), but not in the mutant strain DM103 (*ittA*::Str^R^), when either Northern blot (Fig. 3A) or RT-PCR analysis was employed (Fig. 3B).

**Figure 3.**
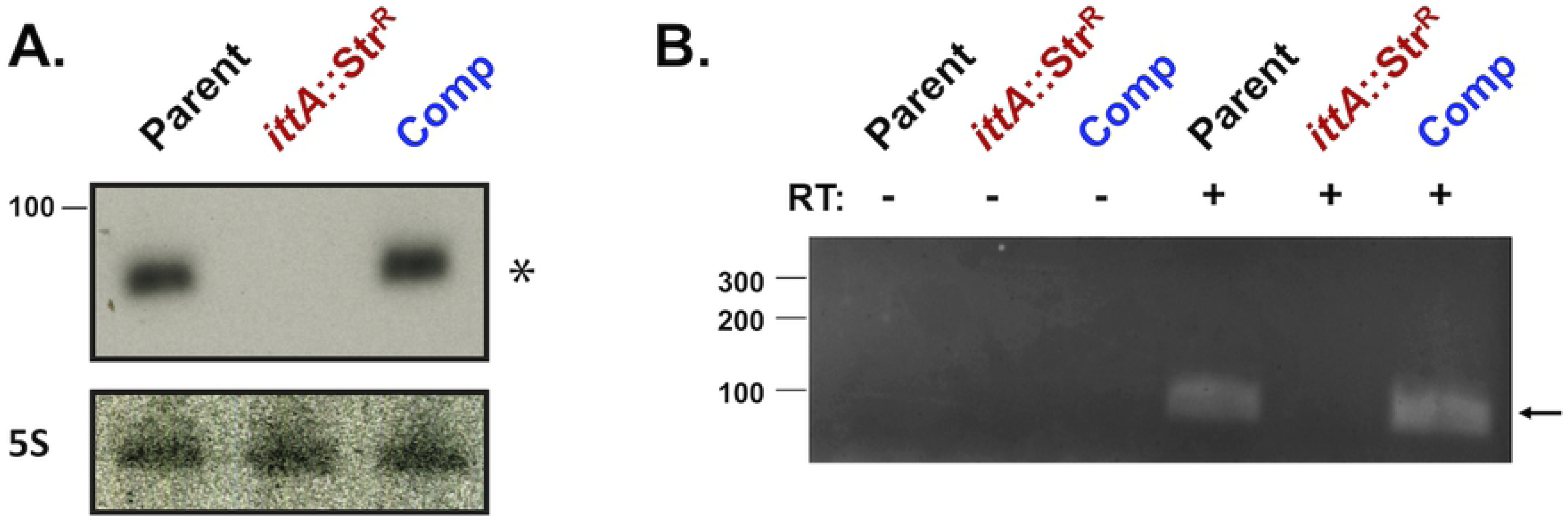
Confirmation that the *ittA* is not made in the mutant strain and is restored in the genetic complement. A. Northern blot of total RNA isolated from the parent strain ML23 (denoted Parent), the sRNA mutant strain (*ittA*:Str^R^), and the *ittA cis* complemented strain (denoted Comp). Detection of 5S from each strain serves as a loading control. B. RT-PCR using purified total RNA from the *B. burgdorferi* ML23 (Parent), sRNA mutant (*ittA*:Str^R^), and genetic complement (Comp) strains. The first three lanes had no reverse transcriptase (RT) added to the reactions (indicated with a “-“) whereas the next three lanes included reverse transcriptase (designated with a “+”). The DNA ladder is shown to the left and base pair values are indicated. The arrow on the right indicates the presence of the sRNA species observed in the parent and complement strains.

Considering the nature of intergenic sRNAs (i.e., between two annotated genes), one concern in deleting the *ittA* sRNA is the potential polar effect this alteration might have on expression of flanking genes. Here, we focused on the only intact encoding genes in the region, *bbd18* and *bbd21.* When RT-PCR was employed, no qualitative difference in *bbd18* or *bbd21* transcripts were observed between the parent, mutant, or complement strains (Fig. S1). Note that both *bbd19* or *bbd20* are no longer annotated as intact ORFs and, consistent with this, no transcript was detected for *bbd20* by RT-PCR for any of the strains (data not shown). Taken together, these data indicate that the inactivation of the *ittA* sRNA exhibits no polar effects. In addition, the ML23 parent, the *ittA* mutant, and the *cis* complement strains all grew similarly, indicating that the loss of the *ittA* sRNA did not impair replication of these borrelial strains under *in vitro* growth conditions (Fig. S2).

### The loss of the *ittA* attenuates *B. burgdorferi* infectivity

We then evaluated the loss of the *ittA*-encoded sRNA on murine infectivity. To spatially and temporally track infection, C3H/HeN mice were inoculated at a 10^3^ dose with the parent *B. burgdorferi* strain, the *ittA* mutant, and the complement. Light emission was quantified at the time points indicated in Fig. 4. As a background control for luminescence, a single infected mouse was not given the luciferase substrate D-luciferin (leftmost mouse in each panel, Fig. 4A). At the dose tested (10^3^ *B. burgdorferi* cells), no signal is detected for any of the strains prior to day 4. At day 4, a clear signal is observed in mice infected with all three strains except for one mouse infected with the sRNA mutant strain (Fig. 4A). Subsequently, the signal increased with a peak at day 7 and then decreased concomitant with the development of the adaptive immune response (Fig. 4A; [64]). All strains displayed similar light emission until day 10 when the *ittA*::Str^R^ strain exhibited a reduction in signal and retained lower light emission through 21 days, e.g., the duration of the infectivity analysis (Fig. 4A). Consistent with the images obtained, quantification of *in vivo* luminescence from the mice revealed significantly lower light emission by the *ittA*::Str^R^ strain compared to the parent on days 10 and on day 14 of infection (Fig. 4B). The complement strain DM113 emitted light comparable to the parent strain on days 7, 14 and 21 (Fig. 4B). These results suggest that the *ittA* complement strain DM113 displays complete *in vivo* complementation during experimental infection (Fig. 4).

**Figure 4.**
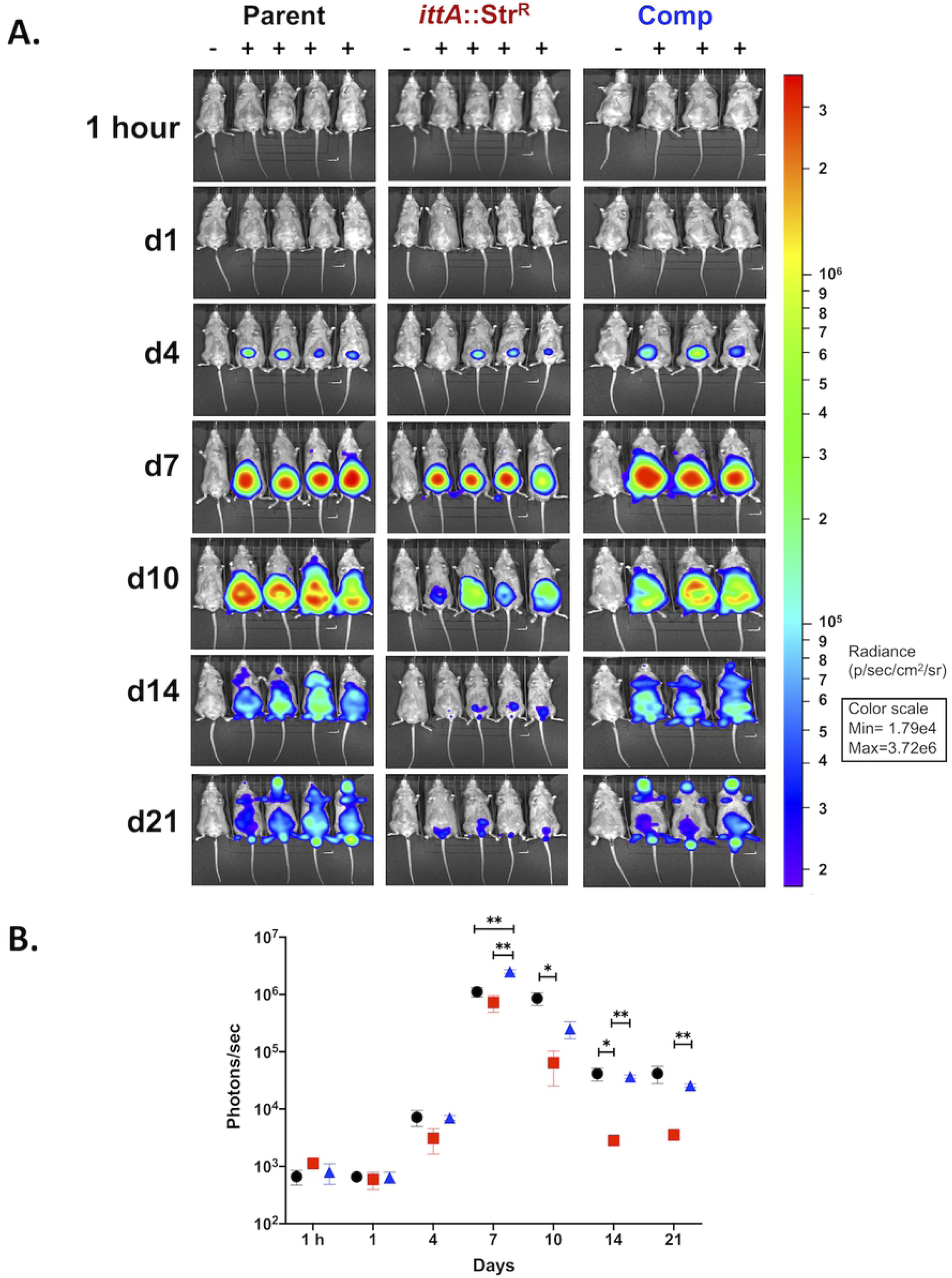
Spatial and temporal infectivity analysis of the *ittA* mutant. A. The course of infection of bioluminescent *B. burgdorferi* strains was tracked following the infection of C3H/HeN mice with 10^3^ of each *B. burgdorferi* isolate. Mice were infected for 21 days total with the parent (n=5), *ittA* sRNA mutant (n=5), and the *ittA* genetic complement (n=4) and imaged on the time or day (d) listed on the left. For each image shown, the mouse on the far left (denoted with a ‘−’) was infected with *B. burgdorferi* but did not receive D-luciferin to serve as a background control. Mice denoted with a ‘+’ were infected with the strain indicated and treated with D-luciferin to promote light emission. All images were normalized to the same scale (shown on the right). B. Quantification of *in vivo* luminescence images of mice infected at a dose of 10^3^ of *B. burgdorferi.* Parent strain ML23/pBBE22*luc* is depicted as black circles, the *ittA* sRNA mutant DM103/pBBE22*luc* as red squares, and the genetic complement strain DM113/pBBE22*luc* as blue triangles. Each time point represents the average value and the standard error from the four mice given D-luciferin substrate for the parent and sRNA mutant strains and three mice for the complement strain. \**p* < 0.05*; **p* < 0.01.

To further assess the phenotype of the *ittA* mutant and complement, the infected mice were sacrificed after 21 days and tissues cultured to qualitatively score for infection. As shown in Table 3, the *ittA* mutant that was needle inoculated on the ventral side (abdomen) is impaired in its ability to disseminate to peripheral ear skin and it is attenuated in its ability to disseminate and colonize heart tissue with infectivity reduced by 40%. In addition to this qualitative assessment of infection, we also scored for spirochete load using quantitative PCR (qPCR) analysis to enumerate borrelial genome copies relative to murine β-actin copies. As observed in Fig. 5, the *ittA* mutant had significantly lower bacterial burden in all tissues analyzed relative to its parent and complemented strains with the notable exception of the inguinal lymph node. The complemented strain exhibited bacterial burden comparable to the parent strain, thus demonstrating complete *in vivo* complementation during infection. When taken together with the *in vivo* imaging data shown in Fig. 4, these data suggest that the *ittA* sRNA is required for optimal tissue tropism and/or dissemination during *B. burgdorferi* experimental infection.

**Figure 5.**
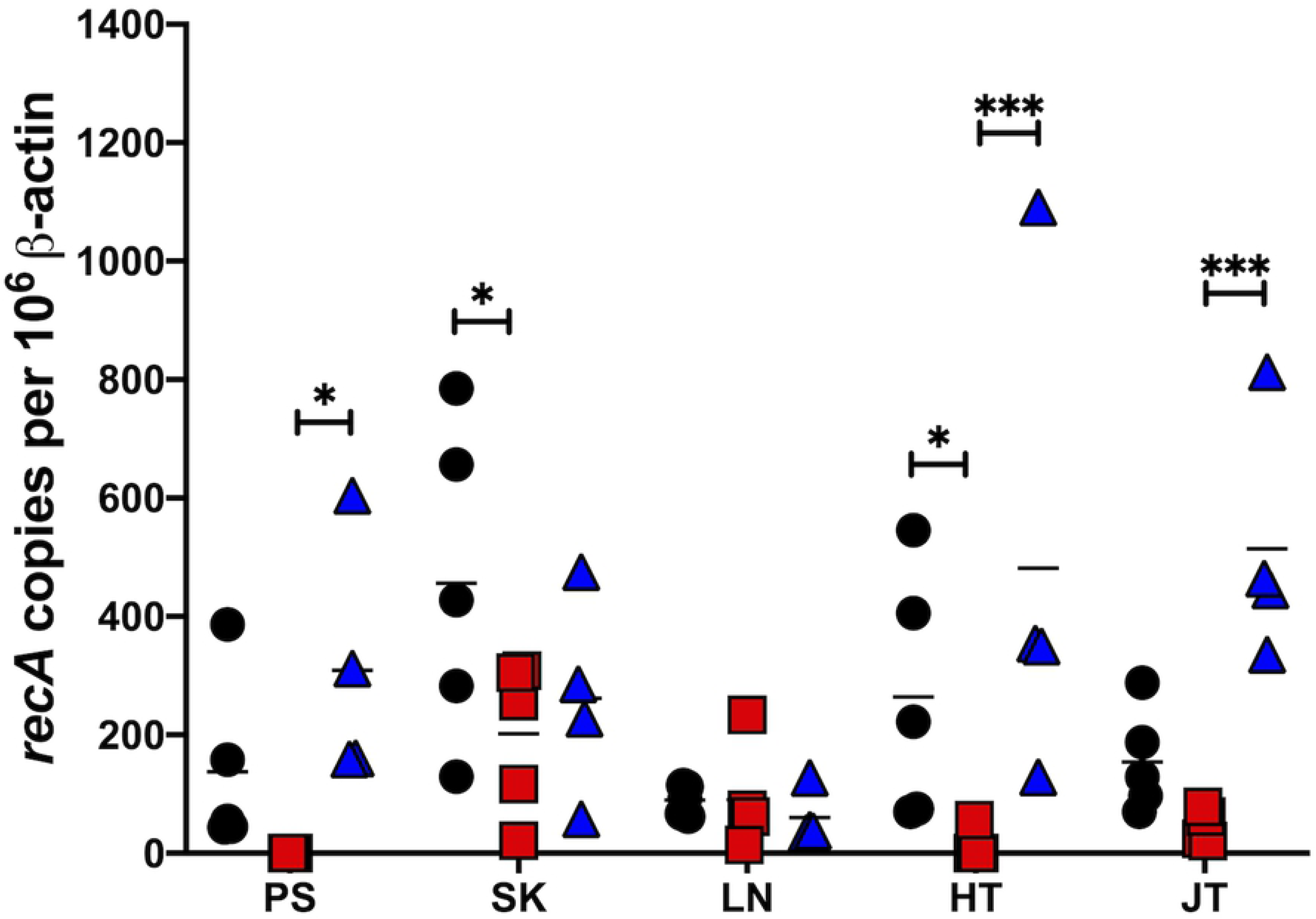
Quantitative assessment of *B. burgdorferi* load from infected mouse tissues. Quantitative PCR (qPCR) of tissues from mice infected with the parental strain (black circles; n=5), the *ittA* mutant (red squares; n=5), and the genetic complement (blue triangles; n=4) was used to enumerate borrelial genomic equivalents relative to the murine samples. Mice were infected with 10^3^ dose of *B. burgdorferi* strains for 21 days. Tissues tested are shown at the bottom: PS for peripheral ear skin; SK for abdomen skin at the site of infection; LN for Lymph node; HT for heart; and JT for the tibiotarsal joint. The results are represented as the number of borrelial *recA* genomic copies per 10^6^ mouse β-actin copies. The horizontal line in each data set depicts the mean value. Each data point shown represents an independent sample from a single mouse tissue assayed in triplicate and averaged. *\* p < 0.05; *** p < 0.001*.

### Transcriptional profile of the *ittA* mutant

*Trans*-acting sRNAs often regulate gene expression by base-pairing with target mRNAs affecting their stability and/or translation. Given the infectivity defect observed, we hypothesized that *ittA* regulates the expression of gene(s) that are important for infectivity and, specifically, skin and heart tissue colonization. We performed RNA-seq to compare the global transcriptional profile of the parent and the *ittA* mutant in an unbiased manner under *in vitro* conditions that mimic mammalian-like conditions. It is important to note that the cultures used for this analysis were also utilized for the subsequent proteomic analysis described below.

In our transcriptional comparison we detected 1,343 transcripts total (Table S1). From this group, we found 92 transcripts that exhibited more than +/- 1.4-fold change and were statistically significant (P_adj_ <0.05; Table S2). Further evaluation showed that the *ittA* mutant exhibited statistically significant, 2-fold change expression of 19 transcripts when compared to the parent (Fig. 6; shown as red spots; see Table S3). Of the 19, 13 transcripts were upregulated in the sRNA mutant strain, including transcripts predicted to be involved in pathogenesis (*vraA* [*bbi16*] [65]), as well as *bba66,* a locus required for effective transmission from the tick vector to mice and expressed during mammalian infection [66, 67]. In addition, there were 6 transcripts downregulated in the sRNA mutant strain, including *ospD* (*bbj09*)*, ospA* (*bba15*), and *oms28* (*bba74*) (Fig. 6). It is important to note that 73 additional genes showed statistically significance (P_adj_ <0.05) with a fold change range between +/-1.4 and +/-1.9 (Fig. 6; shown as gold spots; see Table S4). It is interesting to note that 64 of the significantly affected transcripts are from genes that are known to be BosR/RpoS regulated and 79.7% of them are upregulated in the *ittA* mutant. Overall, these results suggest that the *ittA* sRNA exerts a regulatory effect by either destabilizing or stabilizing several target transcripts or by indirectly altering the expression of these transcripts via an unknown regulatory mechanism. The net effect in the mutant cells lacking *ittA* is a dysregulation of several genes that may contribute to the decrease in fitness and in a reduction in infectivity potential observed.

**Figure 6.**
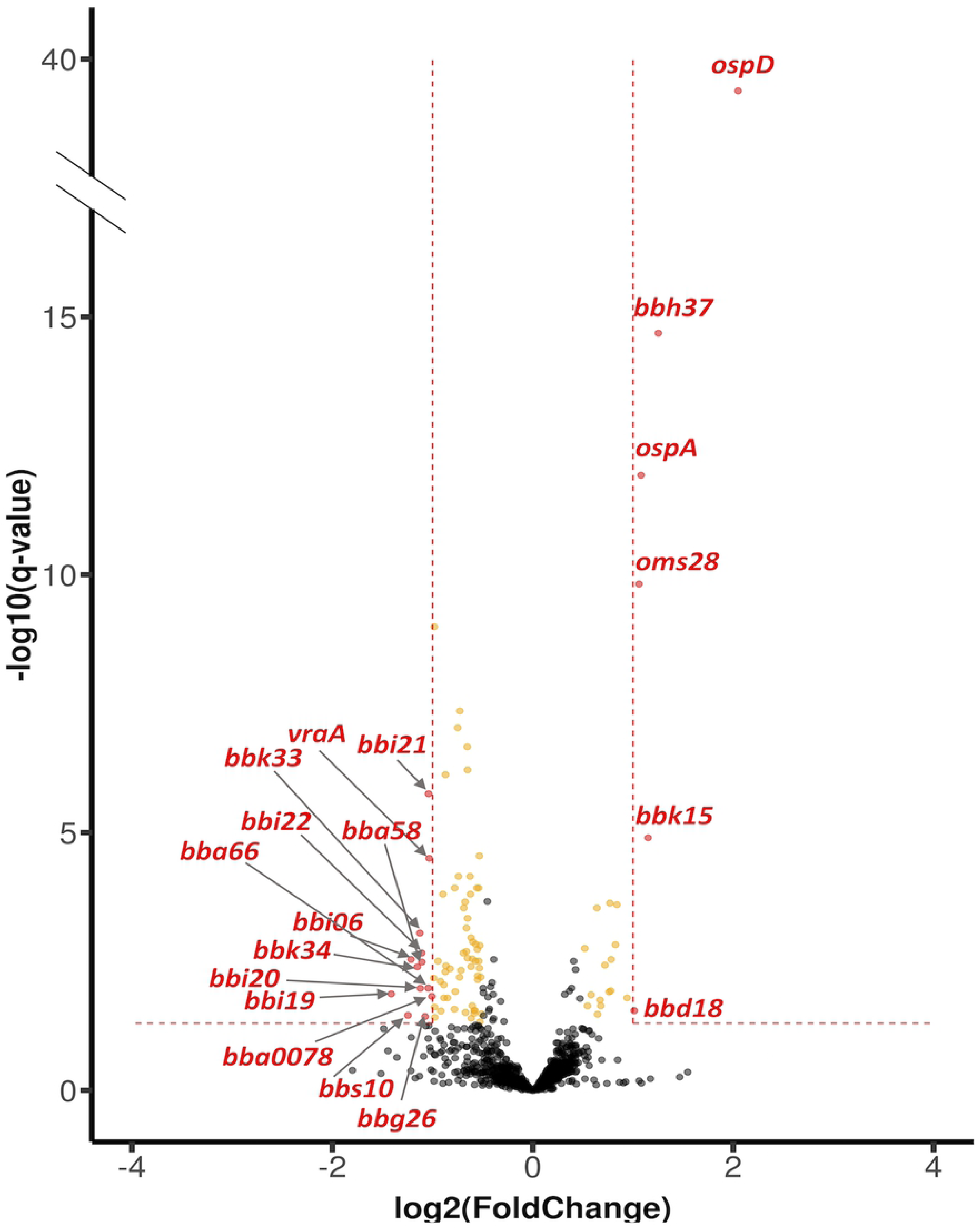
Differential gene expression in the parent relative to the *ittA* mutant. The volcano plot depicts log_2_ fold change on the *x*-axis and False Discovery Rate adjusted *p* value (q -value) on the *y*-axis. The parent and *ittA* mutant strain were grown in biological triplicates *in vitro* using conditions that induce proteins important for mammalian infection. RNA was purified, and the samples subjected to RNA-seq. Single genes are depicted as dots. Of the transcripts, 19 achieved significance with *p*-value <0.05 and with greater than 2-fold change in the mutant relative to the parent strain. Of the differentially expressed genes, 13 were downregulated and 6 upregulated in the parent relative to the mutant. Yellow spots (73 total) are transcripts that achieved significance with a *p*-value <0.05 and a fold change range between +/- 1.4 to 1.9.

### qRT-PCR confirms differential expression of transcripts in the *ittA* mutant

We next utilized quantitative RT-PCR (qRT-PCR) to assess the expression of the candidate genes identified to validate our RNA-seq analysis (Fig. 6) using the parent, *ittA* mutant, and complement strains grown *in vitro* under mammalian-like conditions. A constitutively expressed gene, *flaB*, which was not affected by any of strains tested in this study, was used for normalization [17, 18]. As a control, we tested *ospC* and found that this transcript was not affected by the loss of *ittA*, as predicted. Five genes were tested, *bba66*, *vraA*, *oms28* (*bba74*), *ospD*, and *ospA* (Fig. 7). The qRT-PCR confirmed the upregulation or downregulation observed in the RNA-seq experiment for each gene (Fig. 6 and 7) for the parent and *ittA* mutant. Unexpectedly, only *bba66* was restored to wild type levels in the complemented strain (Fig. 7). Notably, *bbd18* was one of the 19 genes that were differentially regulated. Our qualitative assay demonstrated *bbd18* was expressed in the mutant strain, but our RNA-seq data suggest it may be downregulated in the *ittA* mutant strain. We also attempted to quantify *bbd18* transcripts by both qRT-PCR and Northern blot analysis but were not successful using *in vitro* grown *B. burgdorferi.* Therefore, it is unclear whether *bbd18* expression is altered due to the *ittA* strain construction or *ittA*-dependent gene regulation.

**Figure 7.**
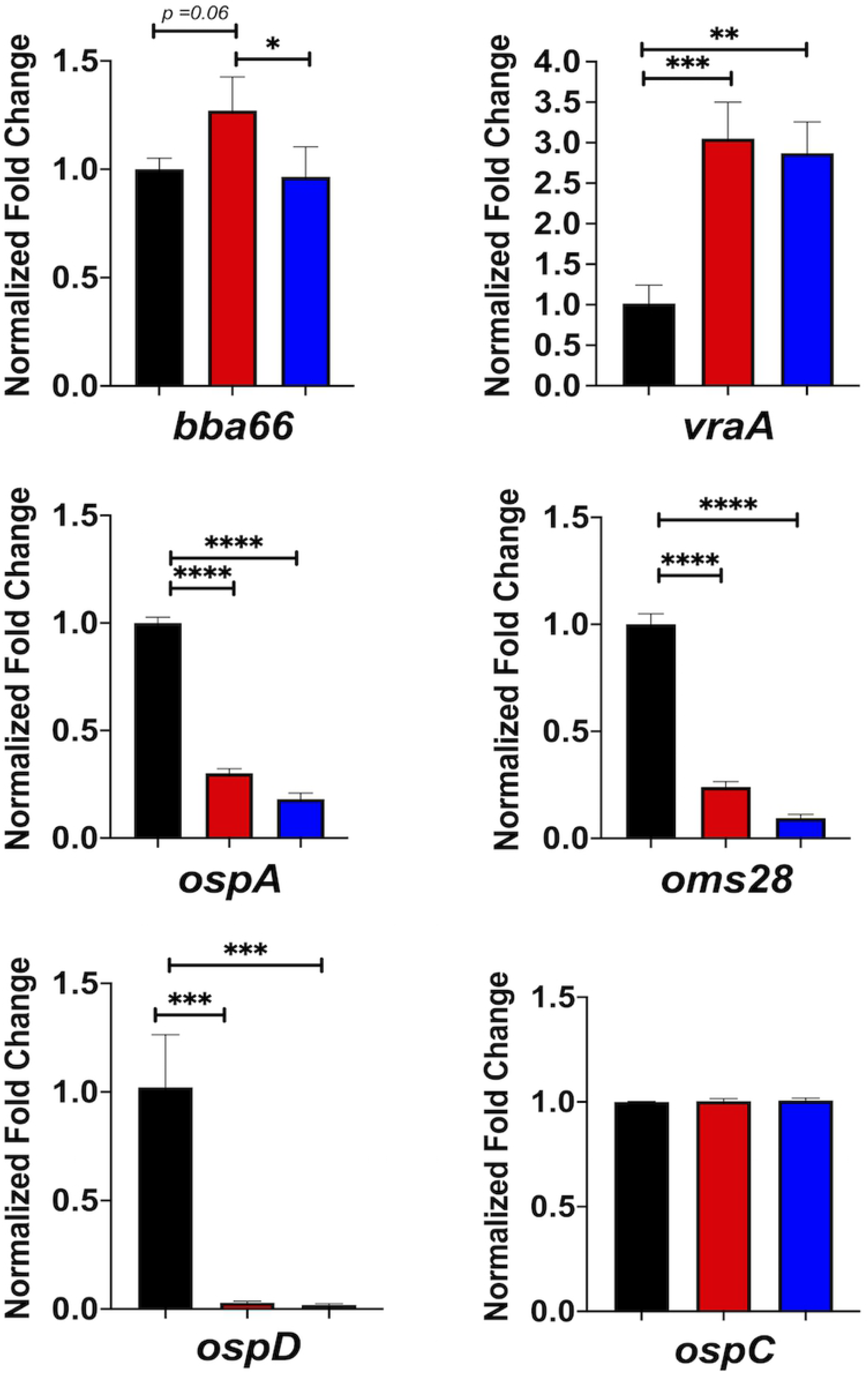
Quantification of transcripts by qRT-PCR. Quantitative RT-PCR (qRT-PCR) of the parent (black bars), *ittA* mutant (red bars), and complemented strain (blue bars) were performed for a subset of transcripts indicated at the bottom of each panel. The data for all samples were normalized to the endogenous control, *flaB,* whose transcription was not affected by the conditions used in the experiment. The error bars indicate standard error. Significance is denotated as * *p < 0.05, ** p < 0.01, *** p < 0.001. **** p < 0.0001*

A limitation in the RNA expression data is that expression of genes differentially expressed in the *ittA* mutant was not restored in the complemented mutant. It is possible that *ittA* is not processed at native levels in the complemented strain. The Northern blot analysis of *ittA* demonstrated that the major sRNA species is the same size and comparable to steady-state levels relative to the parent strain. However, when the Northern blot was overexposed, several bands unique to the complement sample were detected in addition to the major product (Fig. S3). Notably, several of these larger RNA transcripts differ between the parent and the complement strain, suggesting that these differences may affect the *in vitro* readouts tested. Furthermore, the downregulation of *bbd18* in the *ittA* mutant strain may be affecting gene regulation independently of *ittA*.

### *ittA* alters the borrelial proteome

sRNAs often regulate translation initiation by occluding or releasing the ribosome binding site of transcripts [43,47,68]. In addition, sRNAs can bind within coding regions of transcripts resulting in regulatory effects. We hypothesized that *ittA* might modulate translation efficiency of some transcripts at the post-transcriptional level [43,47,68]. To address this, we used a global proteomic screen to identify and quantify the entire borrelial proteome of both the parent and the *ittA* mutant under conditions that mimic mammalian-like conditions (Fig. 8). A total of 718 proteins were detected and are listed in Table S5. Of the 718 proteins, 637 were classified with False Discovery Rate (FDR) of 1% and 81 with FDR of 5%.

**Figure 8.**
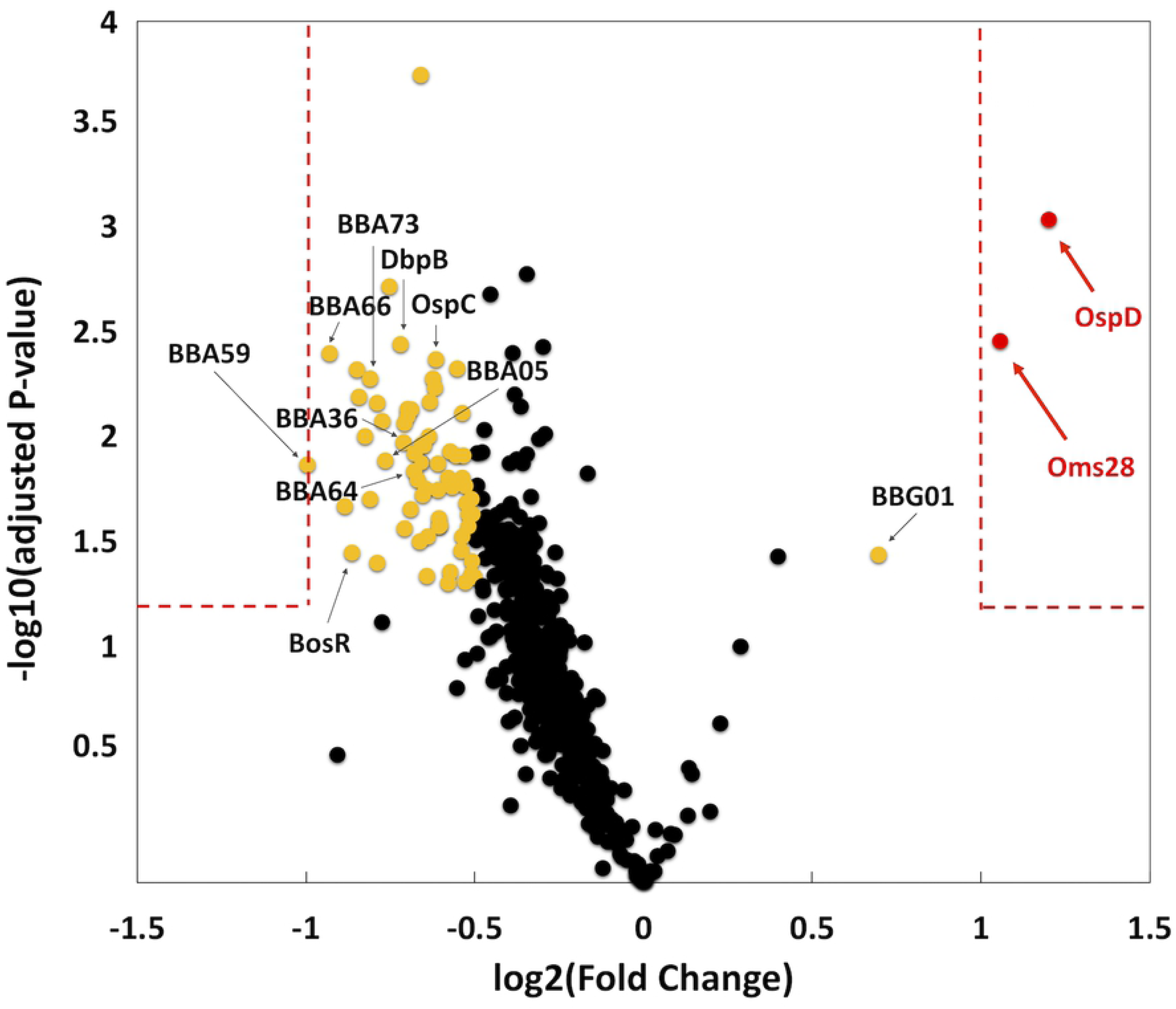
Proteomic evaluation of the parent relative to the *ittA* mutant. Tandem Mass tags (TMT) was used to determine the relative abundance of the total proteome of three biological replicates of parent and *ittA* mutant strains grown under conditions that induce proteins important during mammalian infection. Volcano plot depicts log_2_ fold change (*x*-axis) and log_10_ adjusted *p*-value (*y*-axis) of proteins identified from parent versus the *ittA* sRNA mutant strain. Single proteins are plotted as dots. Proteins outside of the red dashed boxes are significant. Red spots have a +/-2 fold change difference in the parent strain relative to the *ittA* mutant strain and a *p-*value < 0.05. Yellow spots are proteins that achieved significance of *p-*value < 0.05 and a fold change range between +/- 1.4 to 1.9. OspD and Oms28 were found to be significantly higher in abundance in the parent relative to the *ittA* mutant.

The samples used for proteomic analyses were isolated from the same cultures as used for the RNA-seq analysis shown in Fig. 6. From this assessment we found two proteins, OspD and Oms28, that were affected the most by the loss of *ittA*; specifically, they were decreased 2.3- and 2.1-fold, respectively, in the *ittA* mutant (Fig. 8, red spots). In total, 69 proteins were significantly altered by the loss of *ittA*, but 67 of these proteins were in the range of fold change between 1.4- to 1.9-fold (Fig. 8, gold and red spots, respectively, and Table S6). Only one additional protein was made at a statistically higher level in the parent strain relative to the *ittA* mutant (BBG01); all other proteins that fit these criteria were made at higher levels in the *ittA* mutant (Fig. 8 and Table S6). Interestingly, one of the proteins upregulated with fold change of 1.9 in the *ittA* mutant is BBA66, as well as several additional RpoS-regulated surface proteins. Here a subset of RpoS-regulated proteins is synthesized at higher levels in the *ittA* mutant and thus may place the spirochetes at a selective disadvantage since their ectopic production may make them targets for antibody-mediated killing. Presumably this effect is amplified *in vivo* thereby mediating the phenotype observed. However, the complement corrects for this *in vivo* by rescuing the mutant in a manner that is not observed *in vitro*.

Since both the *ospD* and *oms28* transcripts were downregulated in the *ittA* mutant strain in RNA-seq data and proteomic data (Fig. 6 and 8), we hypothesize that *ittA* stabilizes these transcripts and thus allows for the increased translation of OspD and Oms28. We hypothesize that, in the absence of *ittA*, the *ospD* and *oms28* transcripts are destabilized, resulting in reduced levels of OspD and Oms28 proteins. To determine levels of OspD and Oms28 proteins in the *ittA* mutant strain, we performed Western blot analysis of proteins lysates from parent, the *ittA* mutant, and the complement grown *in vitro* under mammalian-like conditions probed with polyclonal antibodies directed against OspD and Oms28. We observed lower levels of OspD and Oms28 in the sRNA mutant than the parent, consistent with the proteomic analysis (Fig. 9). Of note, the complement did not produce OspD to levels similar to that observed in the parent strain, consistent with the expression of *ospD* not being restored in the qRT-PCR analysis (Fig. 7). For Oms28, the complement produced higher proteins levels than the mutant but not to the same level as the parent. These results demonstrate partial complementation of the *ittA* mutant relative to the parent strain using both qRT-PCR and Western immunoblot metrics of assessment.

**Figure 9.**
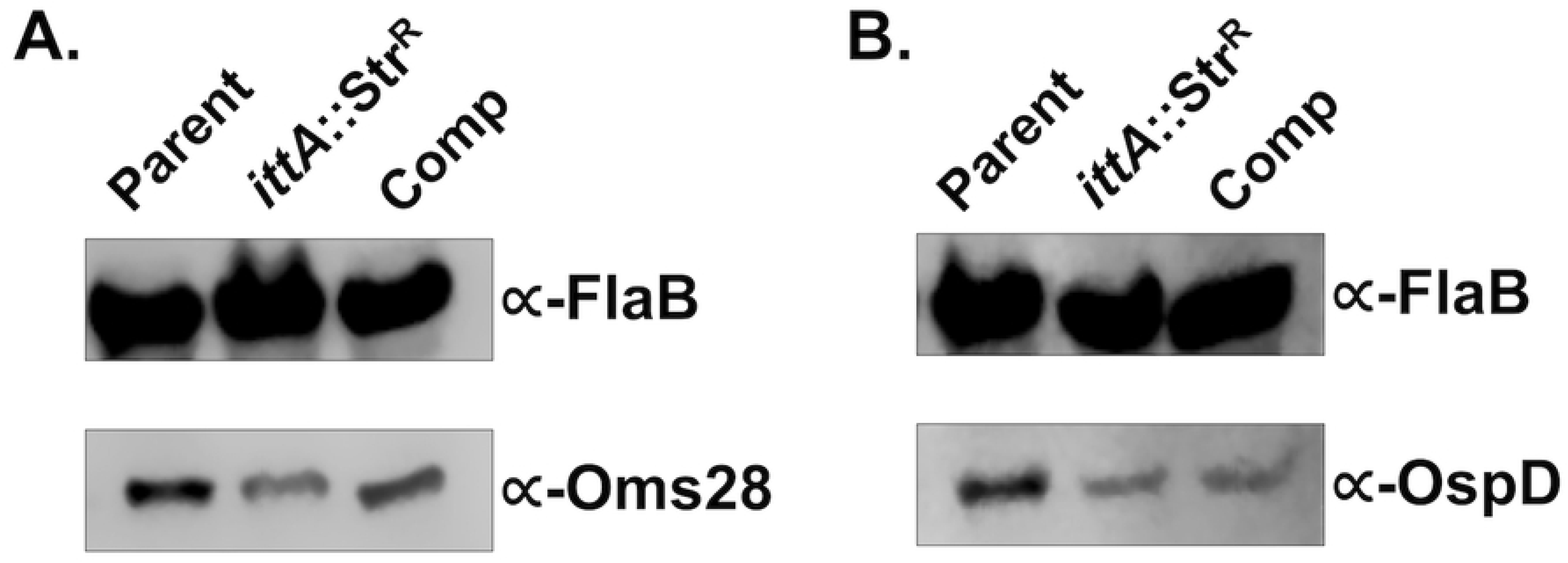
Evaluation of OspD and Oms28 in *B. burgdorferi* lacking *ittA*. Protein lysates were derived from parent, *ittA* sRNA mutant (*ittA*:Str^R^), and complement (Comp) strains grown under conditions that induce proteins important during mammalian infection *in vitro* and probed against anti-OspD (panel A) and anti-Oms28 (panel B). The production of FlaB was used as a loading control for both immunoblots shown.

## DISCUSSION

Small non-coding RNAs (sRNAs) have emerged as an additional mechanism for the regulation of transcript levels or protein function. Specifically, base-pairing sRNAs, such as *trans*-acting sRNAs, bind to transcripts and modulate their expression by either altering their stability or their ability to be translated [47,49,69]. Alternatively, protein-binding sRNAs can bind to cellular proteins and modify their activity [44,45,48]. Regulation via sRNAs is fast acting and functions as an additional layer in response to environmental signals [43, 70]. Despite the significant effect on translation efficiency, it is important to note that the loss of a single sRNA species often has a limited effect on measurable phenotypes. This is likely due to the subtle effect of the sRNA-mRNA interaction and the lack of an absolute effect seen by this event; that is, transcripts are not entirely inhibited or activated depending on the relative abundance of these RNA molecules and the fine-tuning that is associate with the sRNA-mRNA transient interactions [71, 72]. Modest phenotypes may also be due to redundancy, as a given mRNA can be regulated by multiple sRNAs; therefore, the elimination of a single sRNA may not drastically alter mRNA turnover or translation [43,71,72]. Nevertheless, sRNAs encoded by pathogenic bacteria are recognized as important players in adaptive responses with some identified as important effectors in regulatory pathways [43,68,73].

Due to the enzootic nature of *B. burgdorferi* and its ability to quickly adapt to environmental factors encountered during their lifecycle, we hypothesized that *B. burgdorferi* use sRNAs to affect post-transcriptional regulatory processes that calibrate these responses. Recently, 1,005 sRNAs were identified in *B. burgdorferi*, suggesting that these spirochetes exploit this type of genetic regulation [40]. However, how *B. burgdorferi* utilizes these sRNA candidates remains largely unknown. Understanding the role these sRNAs play in post-transcriptional gene regulation in *B. burgdorferi* should provide significant insight into how the Lyme disease spirochete refines its molecular pattern to survive and persist within the disparate environments that they reside, e.g., arthropods and mammals. In this study, we inactivated a *trans*-acting, intergenic (IG) sRNA, designated *ittA*, and demonstrated that this sRNA is required for optimal infection, as well as dissemination to and/or survival in distal tissues. It was further shown that transcript and protein production are affected when the *ittA*-encoded sRNA is not expressed in *B. burgdorferi*. To our knowledge, no other intergenic sRNA from *B. burgdorferi* has been characterized to this extent.

As a first step to link a *B. burgdorferi trans*-acting intergenic sRNA to an infectivity phenotype, mice were infected with the transposon library of *B. burgdorferi* strain B31 [54] and decreased infectivity was scored using Tn-seq [55]. These data were compared against the recently described *B. burgdorferi* sRNA-specific library to identify intergenic sRNA species that mapped to existing Tn mutants. We tested each identified intergenic sRNA Tn mutant individually using the experimental murine infection model. One sRNA mutant, initially described as SR0736 [40] and renamed *ittA* herein, exhibited the most severe infectivity defect. Subsequent *in vivo* imaging of an independently derived mutant in *ittA* confirmed the attenuated phenotype observed for the qualitative infectivity assessment (Table 2) relative to the parent and complement (Fig. 4). The inactivation of *ittA* affected tissue tropism of *B. burgdorferi* where spirochetes were cultured out of peripheral ear skin and heart in less than 8% of the samples tested (Table 2). Even though the qualitative assessment of infection showed some degree of colonization in the remaining tissues (except the lymph node), the total bacterial load of *B. burgdorferi* was significantly lower for the *ittA* mutant relative to the parent and complement strains (Fig. 5). These data indicate that *ittA* is needed for optimal borrelial colonization and/or dissemination. Furthermore, it is important to note that the *ittA* complement strain DM113 completely restored infectivity in a manner indistinguishable from the parent using infection as our readout, indicating that the phenotype of the *ittA* mutant was due to its absence and not a second site mutation.

**TABLE 2.** Infectivity of *B. burgdorferi* intergenic (IG) sRNA transposon mutants relative to their genetic parent clone 5A18NP1^a^

| Strain<sup>b</sup> | sRNA ID<sup>c</sup> | Genomic location | Ear | Skin<sup>d</sup> | Lymph node | Heart | Bladder | Joint | Total sites | % Positive Tissues |
| --- | --- | --- | --- | --- | --- | --- | --- | --- | --- | --- |
| **5A18NP1 (parent)** | N/A | N/A | 4/4 | 4/4 | 4/4 | 4/4 | 4/4 | 4/4 | 24/24 | 100% |
| **T05TC355** | SR0897 | lp54; IG *bba34-bba36* | 3/4 | 3/4 | 3/4 | 3/4 | 3/4 | 3/4 | 18/24 | 75% |
| **T10TC061** | SR0912 | lp54; IG *bba66-bba68* | 4/4 | 4/4 | 4/4 | 4/4 | 4/4 | 4/4 | 24/24 | 100% |
| **T11TC387** | SR0948 | cp26; IG *bbb03-bbb04* | 4/4 | 4/4 | 4/4 | 4/4 | 4/4 | 4/4 | 24/24 | 100% |
| **T10TC351** | SR0962 | cp26; IG *bbb13-bbb14* | 2/4 | 1/4 | 2/4 | 2/4 | 2/4 | 2/4 | 11/24 | 46% |
| **T04TC273** | SR0725 | lp17; IG *bbd04-bbd05a* | 2/4 | 2/4 | 2/4 | 2/4 | 2/4 | 2/4 | 12/24 | 50% |
| **T06TC412** | SR0736 | lp17; IG *bbd18-bbd21* | 0/4 | 1/4 | 3/4 | 0/4 | 2/4 | 0/4 | 6/24 | 25% |
| **T08TC464** | SR0795 | lp28-3; IG *bbh36a-bbh36b* | 4/4 | 4/4 | 4/4 | 4/4 | 4/4 | 4/4 | 24/24 | 100% |
| **T09TC062** | SR0853 | lp38; IG *bbj15a-bbj16* | 3/4 | 2/4 | 3/4 | 3/4 | 3/4 | 3/4 | 17/24 | 71% |

<sup>a</sup> Dose of 10<sup>4</sup> per strain of *B. burgdorferi* tested; <sup>b</sup> Transposons from Tn library of *B. burgdorferi* [54] interrupting sRNAs; <sup>c</sup> sRNAs from *B. burgdorferi* sRNA library [40]; <sup>d</sup> skin from abdomen (inoculation site).

**TABLE 3.**
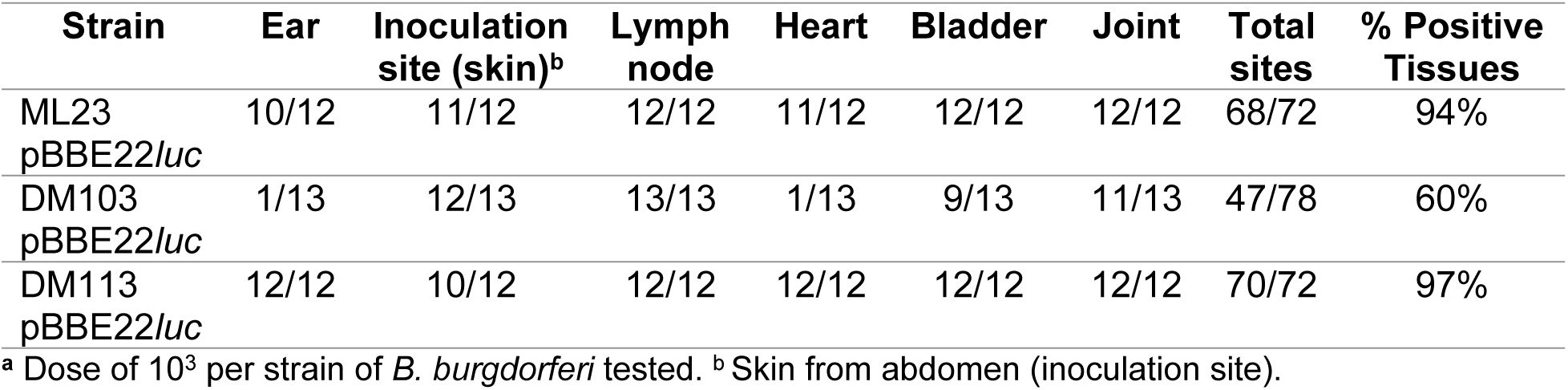
Infectivity of the sRNA mutant strain relative to its parent and genetic complement^a^

To globally assess the role of the *ittA* sRNA we used both transcriptomic and proteomic approaches to identify transcripts and proteins that are altered in its absence, respectively. Two proteins, OspD and Oms28 (BBA74), were produced at lower levels in the *ittA* mutant consistent with their respective transcripts being reduced in the mutant background. We hypothesize that the *ittA* sRNA might bind to the *ospD* and *oms28* transcripts, prevents their degradation, and consequently enhances OspD and Oms28 translation. Whether this sRNA-mRNA interaction occurs and affects the *ospD* and *oms28* transcripts directly, or if this effect is due to other *ittA*-regulated targets that indirectly affect this process, remains to be determined.

We aimed to validate five of the nineteen transcripts identified as being altered due to the loss of the *ittA* sRNA: *ospA*, *ospD*, *oms28*, *vraA*, and *bba66*. Of these five, *bba66* and *vraA* have been associated with some aspect of mammalian-based virulence; four of these genes, including *ospA*, *ospD*, and *oms28*, as well as *bba66*, are expressed in the arthropod vector and some have significant phenotypes in this stage of the *B. burgdorferi* life cycle, particularly *ospA* and *bba66* [66,67,74–80]. OspA is a well characterized surface lipoprotein that functions as an adhesin in the midgut of *Ixodes* ticks [8,76,77,81]. BBA66 is a lp54 encoded surfaced expressed lipoprotein that is upregulated during nymph blood meal and is highly expressed in the mammal for an extended period of time, suggesting a role in persistence [67]. Needle inoculation of mice with *B. burgdorferi bba66* mutants results in lower bacteria burden in joint tissue and significantly lower joint swelling relative to the parent strain, suggesting that BBA66 contributes to borrelial-mediated inflammation [66]. Mutants in *bba66* are acquired by larvae and persist through molting, but were significantly impaired in their ability to infect mice when introduced by tick bite compared to that of mice fed upon by ticks seeded with wild type *B. burgdorferi*, suggesting a role for BBA66 in transmission [66]. Recently, an additional function of BBA66 was found, whereby BBA66 binds to the neuroglial and macrophage protein Daam1 [82]. Daam1 is in the formin family of proteins involved in regulating cytoskeletal reorganization in mammalian cells [83]. A prior study showed that Daam1 co-localized to pseudopods on macrophages that processed *B. burgdorferi* by coiling phagocytosis [84]. A more recent report showed that BBA66 mediated the attachment of *B. burgdorferi* to these cells via the Daam1 protein [82]. Interestingly, the *bba66* mutant exhibited reduced levels of internalized *B. burgdorferi* while borrelial cells that produced greater amounts of BBA66 were phagocytosed more efficiently [82]. In the RNA-seq data shown (Fig. 6), higher expression of *bba66* was observed in the sRNA mutant compared to the parent. In addition, in Fig.8 and Table S6, the mutant has a 1.9 fold change increase in protein abundance of BBA66 relative to the parent (Fig. 8 and Table S6). It is tempting to speculate that the increase of *bba66* expression in the sRNA mutant may lead to increased phagocytosis and local clearance of *B. burgdorferi* by macrophages as a result of BBA66-mediated binding of Daam1. This could help explain the lower bacteria burden in the majority of the mice tissues examined in the sRNA mutant, where BBA66 levels are predicted to be greater (Fig. 4). In Fig. 10, predicted secondary structure of *ittA* and interaction with *bba66* is displayed. Whether this sRNA-mRNA interaction occurs and affects *bba66* transcript directly remains to be determined. Taken together, the dysregulation of *bba66* expression may be an important factor in the phenotype observed for the *ittA* mutant (Fig. 4 and 5).

**Figure 10.**
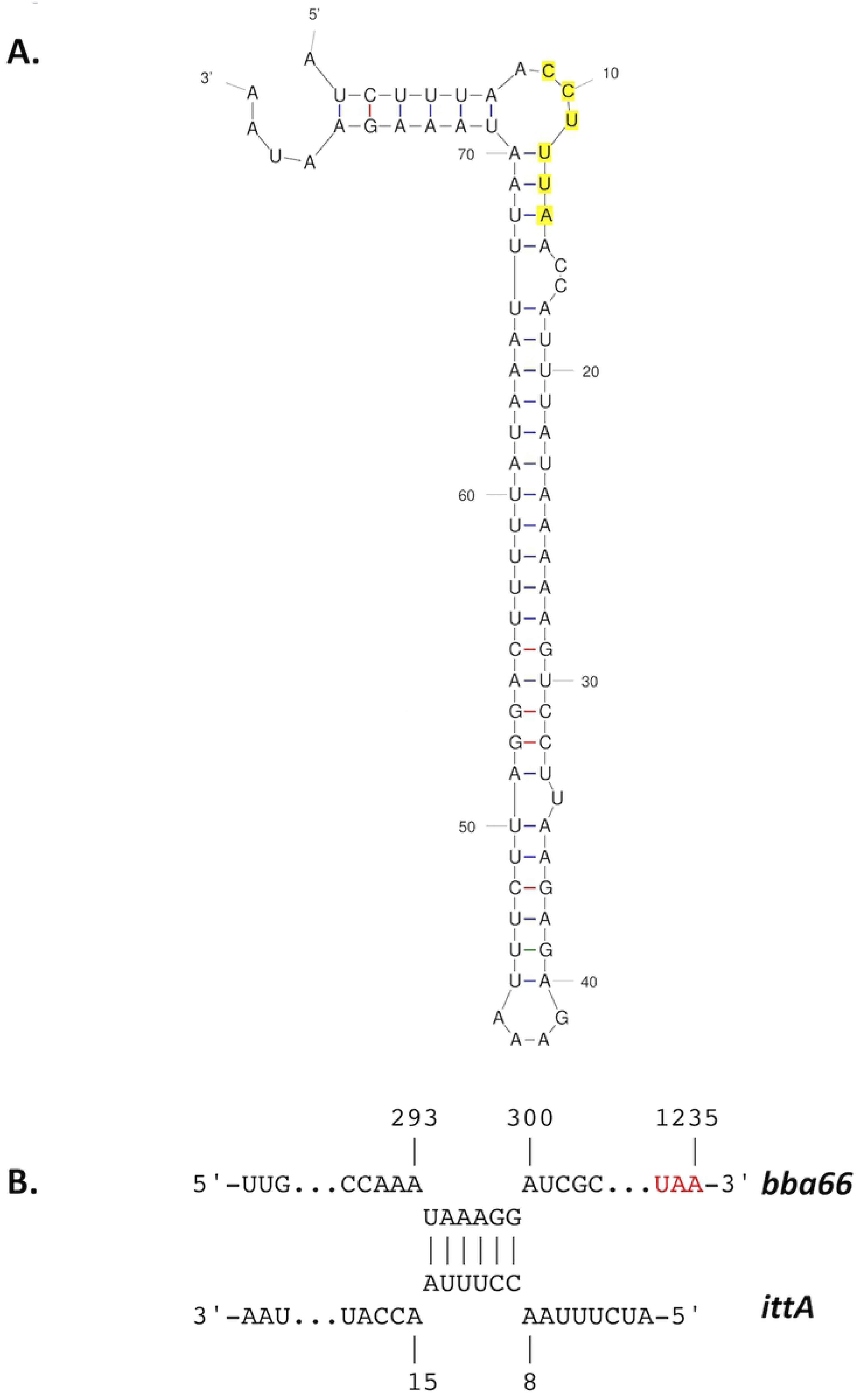
IntaRNA (Freiburg RNA Tools) predictions of interactions of the *ittA* sRNA with the *bba66* transcript. A. A predicted secondary structure of the *ittA* sRNA (mfold). B. Potential interaction of the *ittA* sRNA (lower) within the coding sequence of *bba66* (upper). The red depicts the *bba66* stop codon.

In addition to BBA66, several BosR/RpoS-regulated genes and the proteins they encode are present in the *ittA* mutant *in vitro* at higher levels than the parent (Fig. 6 and 8), suggesting that the loss of this sRNA leads to mis-regulation of surface exposed proteins during mammalian infection. Additional RpoS-regulated targets are subject to mis-regulation in the *ittA* mutant include OspC, DbpA, DbpB, and BBK32, which all contribute to borrelial pathogenesis [60,85–92]. That these virulence-associated proteins are produced more in the mutant relative to the parent suggests that the defect here may be due to ectopic mis-regulation that place the spirochete at a selective disadvantage. Along these lines, previous studies have shown that the mis-regulation of *ospC*, in the form of constitutive expression, results in the clearance of *B. burgdorferi* in experimental infection [85, 86]. It is thus likely that a coordinated mis-production of several of these proteins could have a detrimental synergistic effect that clears *B. burgdorferi* in experimentally infected mice.

One significant and confounding issue stemmed from the incomplete *in vitro* complementation of the *ittA* mutant. Notably, the *cis*-acting complement restored infectivity to wild type levels as assessed by all *in vivo* metrics tested, e.g., *in vivo* imaging as well as both qualitative and quantitative measure of infected tissues, *in vitro* indicators were not fully reinstated. Surprisingly, during *in vitro* growth, several of the transcripts and proteins affected by the absence of the *ittA* sRNA were not restored to wild type levels in the complement. It is possible that factors impacting the gene context for *ittA* (e.g. the presence of the downstream gentamicin cassette) or secondary genetic changes in the complemented strain may affect *ittA* sRNA expression, processing, or stability during *in vitro* culture conditions. The *in vitro* complement strain results indicate some caveat to the mutant or complement strain construction (discussed below). However, the *in vivo* data demonstrate that *ittA* is an important factor for infectivity. As mentioned previously, additional RNA species containing *ittA* sequences were detected by Northern blot analysis in the complemented strain. Additional experimentation is planned to resolve this issue.

Of the five targets tested for transcript levels, only *bba66* was restored to wild type levels in the complemented strain. A predictive algorithm, IntaRNA (Freiburg RNA tools, [93]) indicated potential *ittA* binding site within the *bba66* transcript with the 5’ loop of the predicted *ittA* secondary structure. We hypothesize that the *ittA*-*bba66* RNA interaction may lead to RNase-dependent degradation of the *bba66* transcript (Fig.10), since *bba66* is expressed in higher amounts in the *ittA* mutant than the parent and complement strains. Further study is needed to determine whether this mechanism is active in the case of *bba66* regulation.

At the protein level, we were limited by available antibody reagents and, of the two candidates tested, OspD, and Oms28, only Oms28 appeared partially restored to the levels observed in the parent strain in Western immunoblotting. Although the *cis* complementation strategy was designed to create as little a difference relative to the parent strain as possible, the addition of an additional antibiotic resistance marker may alter the expression of the *ittA* sRNA in a manner that yields an atypical regulatory response. In this regard, the Northern blot shown in Fig. 3, when exposed longer, showed a unique unprocessed form of the *ittA* sRNA (Fig. S3). Whether this contributes to the complementation defect remains to be determined. Another possibility is that the overlap of the initial 5’ transcription start site of *ittA* with the transcriptional start site of *bbd18* on the opposite DNA strand (Fig. 1) results in some interference that affects the ability of *ittA* to carry out its regulatory effect(s) or affects the levels of BBD18. Why this effect is limited to our *in vitro* studies but not observed *in vivo* is not clear. Further studies will be needed to address this experimental conundrum.

In conclusion, sRNA-mediated regulation has been proven to be complex but important in fine-tuning gene expression in many bacteria species [94, 95]. Our findings indicate that the *ittA* sRNA is required for optimal infectivity and that its absence alters the expression and production of a number of genes and proteins, respectively. One possibility posits that *ittA* may help in a quick adaptive response that alters the translation efficiency in a number of transcripts. The effect of each gene regulated by *ittA* individually is subtle but collectively the net effect may result in the dysregulation of these targets, yielding a synergistic response that results in an attenuated phenotype. Here, there is an additional layer of complexity, seen in the form of tissue tropism, such that colonization at remote skin and cardiac tissue sites is impaired. This work thus suggests that sRNA-based regulation via *ittA* is important in maintaining appropriate levels of gene expression that promote *B. burgdorferi* colonization and dissemination during experimental infection.

## MATERIALS AND METHODS

### Bacteria strains and culture conditions

Bacterial strains and plasmids used in this study are described in Table 4. *Escherichia coli* strains were grown aerobically at 37°C in Luria Broth (LB). Concentration of antibiotics used in *E. coli* are as follows: kanamycin, 50 μg/ml; spectinomycin, 50 μg/ml; and gentamicin, 5 μg/ml. *B burgdorferi* strains were grown in BSK-II media supplemented with 6% normal rabbit serum (Pel-Freez Biologicals, Rogers, AR) under conventional microaerobic conditions at 32°C, pH 7.6, under 1% CO_2_ atmosphere, or conditions that induce genes and gene products important in the mammalian environment, namely, 37°C, pH 6.8, under 5% CO_2_ atmosphere. *Borrelia burgdorferi* B31 ML23 [59] and derivative strains were grown under antibiotic selective pressure, dependent on genetic composition, with either kanamycin at 300 μg/ml, streptomycin at 50 μg/ml, or gentamicin at 50 μg/ml. *B. burgdorferi* 5A18NP1 [57] was grown in the presence of kanamycin at 300 μg/ml and the *B. burgdorferi* transposon mutants [54] were grown in the presence of both gentamicin at 50 μg/ml and kanamycin at 300 μg/ml.

**TABLE 4.**
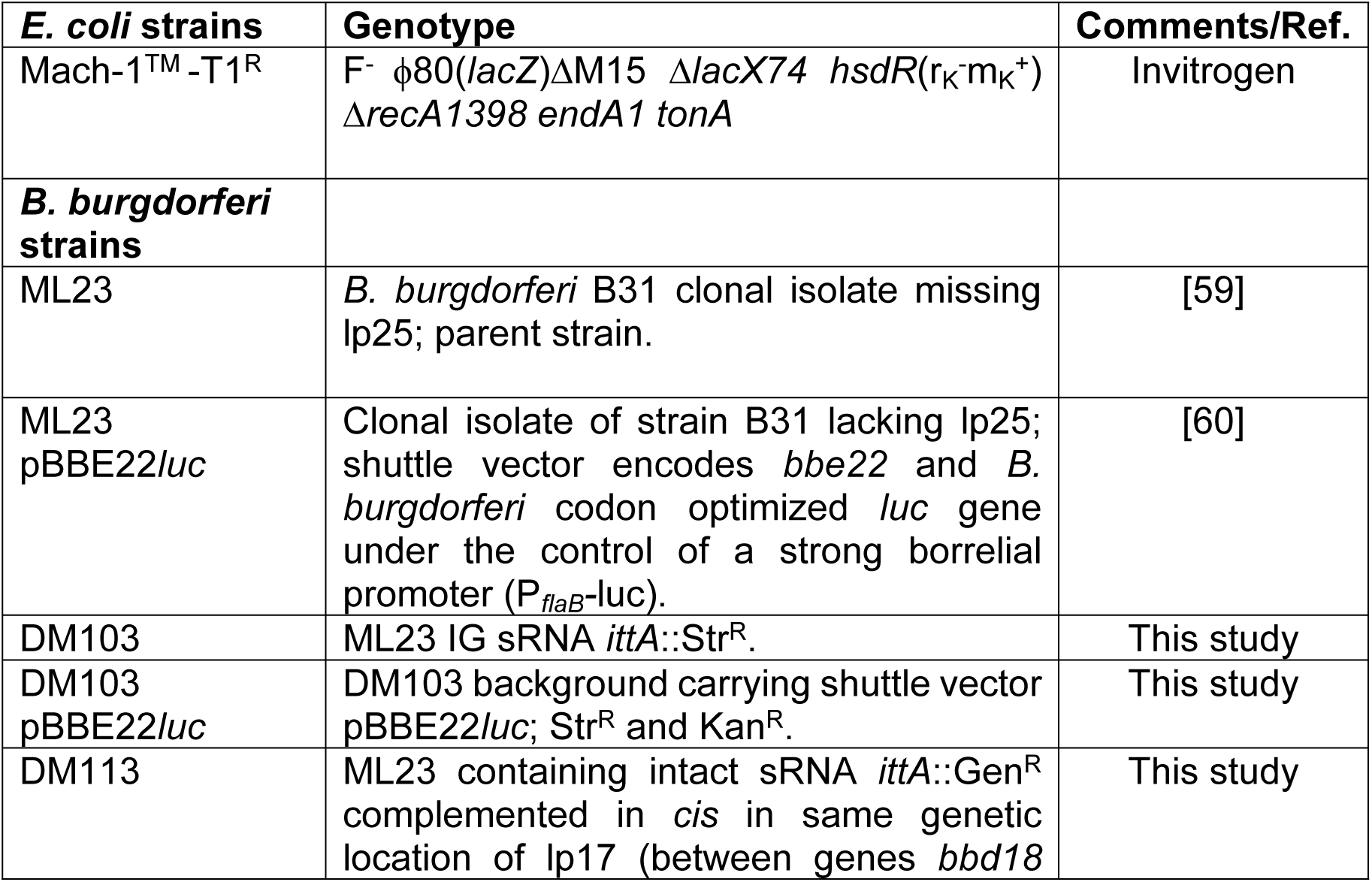
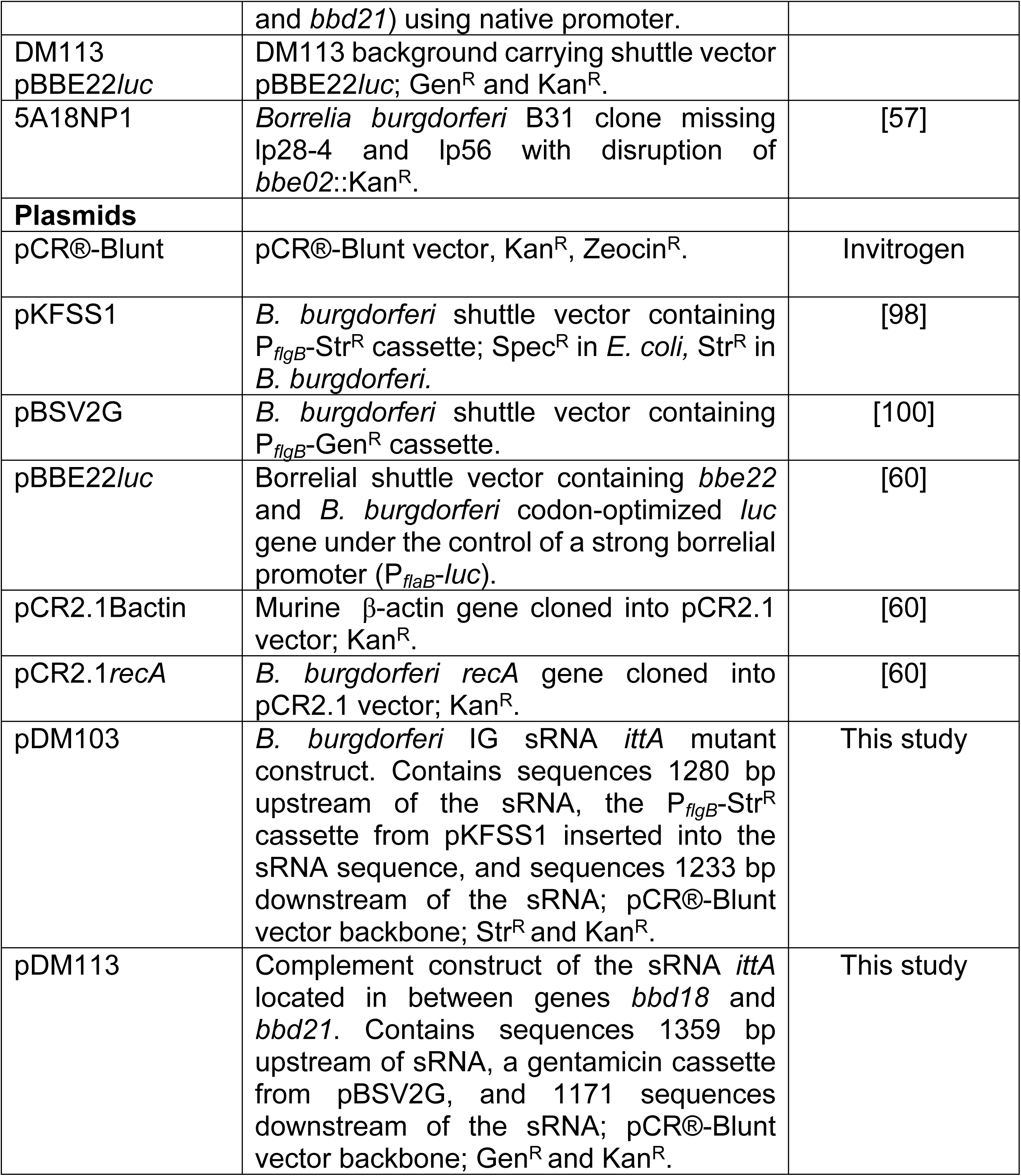
Plasmids and strains used in this study.

### Transposon mutagenesis screen

Transposon mutants were obtained from the arrayed *B. burgdorferi* library [54]. In order to conduct Tn-seq experiments, a single pool containing all of the Tn mutants was generated by combining sub-pools containing 70-80 individual Tn mutants each [54,96,97]. The Tn library was grown for 48 hours in the presence of kanamycin and gentamicin. Cell density was determined by dark-field microscopy. Groups of six 9-14 week old C3H/HeJ mice (Jackson laboratories) were injected in the right flank with 5x10^5^ *B. burgdorferi* by needle inoculation. As a control to prevent *in vitro* growth defects from affecting the results of the *in vivo* screen, 5x10^5^ organisms from the inoculum used to inject the mice were cultured in 12 ml BSK-II broth supplemented with kanamycin and gentamicin. The *in vitro* control cultures were grown for 3 days to parallel time in culture for the tissue cultures. The bacteria in the culture were collected using centrifugation for 20 minutes at 3,000 x *g* and the pellet frozen at -80°C.

The mice infected with the transposon library were sacrificed two weeks post-infection. The tibiotarsal joint closer to the inoculation site was removed under aseptic conditions. The tibiotarsal joints from each group of injected mice were cultured together in 12 ml of BSK-II broth supplemented with kanamycin and gentamicin. The cultures were checked daily for growth. When the density of the cultures reached late exponential phase, the bacteria were centrifuged and the pellet frozen as described above.

Genomic DNA was obtained from the frozen bacteria pellets using a DNeasy Blood and Tissue Kit (Qiagen, Valencia, CA) as per the manufacturer’s instructions. For the *in vivo* samples, genomic DNA obtained from the organ cultures of different groups of infected mice were pooled in relative proportions such that each library for sequencing represented the bacteria recovered from 24 mice. For the *in vitro* control samples, DNA recovered from the cultures of the inoculums were pooled to match the groups of mice pooled to make each of the *in vivo* sample libraries. Libraries for sequencing were constructed as described previously [97]. Briefly, an aliquot of the genomic DNA was placed in a 2 ml microfuge tube and sheared by sonication. Cytosine tails (C-tails) were added to 1 µg sheared DNA using terminal deoxynucleotidyl transferase (TdT, Promega, Madison, WI). Transposon containing fragments were amplified in a PCR containing DNA from the TdT reaction as template and primers specific to the ColE1 site on the 5’ end of the transposon, pMargent1 and the C-tail, olj376 (Table S7). To prepare the DNA for sequencing and further amplify the transposon-genomic DNA junction, a nested PCR reaction was performed using DNA from the first PCR as a template, a primer specific to the transposon end, pMargent2 (Table S7), and an indexing primer containing the specific sequences required for sequencing on an Illumina platform and where NNNNNN represents a six-base-pair barcode sequence allowing samples to be multiplexed in a single sequencing lane. Within an experiment, a unique indexing primer was used for each individual *B. burgdorferi* sample. A majority of the PCR products were between 200 and 600 bp. The sequencing libraries made from the *in vitro* and *in vivo* samples were pooled at equal concentrations prior to sequencing. The pooled libraries were sequenced on an Illumina HiSeq 2500 at the Tufts University Core Facility as 50 bp single-end reads using the custom sequencing primer pMargent3 and the standard Illumina index primer.

Sequence data analysis was performed as described previously [97]. Briefly, sequenced reads were aligned to the *B. burgdorferi* B31 genome using the short read aligner Bowtie. A custom script was then used to compile the resulting SAM files into a Microsoft Excel spreadsheet with the number of reads aligned to each site or annotated gene listed. The frequency of transposon mutants with insertions in a particular site or gene in the population was assessed by determining the number of sequence reads aligned to that mutant or gene as a percentage of all reads in a given sample.

### Genetic inactivation of the *B. burgdorferi ittA* intergenic sRNA

The intergenic (IG) small regulatory RNA (sRNA) located between genes *bbd18* and *bbd21* in lp17 was insertionally inactivated via homologous recombination by replacing the 3’ end of the sRNA (nucleotides 11,820-11,850) with the P*_flgB_*-*aadA* (streptomycin resistant; Str^R^) antibiotic cassette [98] . This sRNA was designated SR0736 by Popitsch *et al*. [40]. Based on the data obtained herein, we renamed the SR0736 sRNA *ittA*. DNA sequences that flanked the *ittA* sRNA locus were amplified using PCR with PrimeSTAR GXL polymerase (Takara, Mountain View, CA). For the upstream fragment, 1280 bp were amplified using primers US-F and US-SpecR (see Table S7). The 1266 bp fragment containing P*_flgB_*-Str^R^ was PCR amplified from pKFSS1 [98] using the oligonucleotide primers pair US-SpecF and SpecDS-R (Table S7). An additional 1,233 bp PCR product, which amplified sequences downstream from the *ittA* sRNA, was engineered with primers SpecDS-F and DS-R (Table S7). All three fragments had 20 base pair overlap sequences and were assembled by overlap PCR [62,63,99].

For the creation of the sRNA *cis* complement strain, the P*_flgB_*-Str^R^ cassette of the sRNA mutant was replaced on lp17 using the native *ittA*-containing sequence with a linked P*_flgB_*- Gent^R^ marker downstream of the sRNA [100]. The US-F and compUS-gentR primers were used to PCR amplify the 1359 bp portion containing *ittA* (Table S7). The 983 bp gentamicin cassette from pBSV2G [100] was produced using primers compUS-gentF and compgentDS-R (Table S7). A 1171 bp region downstream from *ittA* was amplified using the primers compgent-DSF and compDS-R (Table S7). As before, all three fragments had 20 base pair overlap sequences and were assembled by NEBuilder HiFi DNA Assembly Master Mix (New England Biolabs, Ipswich, MA). All constructs were verified by Sanger sequencing prior to transformation into the *B. burgdorferi* strain B31 derivative ML23 [59] or DM103.

### Transformation of *B. burgdorferi*

*B. burgdorferi* were made competent for DNA transformation as previously described [62,63,90,101]. Prior to transformation via electroporation, all plasmid constructs were linearized with *Xho*I. Following antibiotic selection for the desired strain, putative transformants were tested for the presence of the genetic constructs using a PCR-based screen using primers from Table S7. Subsequent mutant or complemented strains were tested for *B. burgdorferi* strain B31 plasmid content by PCR [18].

### Infectivity studies and bioluminescent imaging

Infectivity studies were performed as previously described [60,62,63,90]. Briefly, 8-week-old C3H/HeN female mice were inoculated with 10^3^ organisms of the *B. burgdorferi* parent strain ML23/pBBE22*luc,* the sRNA *ittA*::Str^R^ strain DM103/pBBE22*luc,* or the genetic complement strain DM113/pBBE22*luc,* by intradermal injection in the abdomen skin. For the parent and complement strains, twelve mice were infected, for the sRNA inactivation strain, thirteen mice were infected.

The bioluminescent imaging was performed as done previously [60,62,63]. Briefly, five mice were imaged for the parent and mutant strains and four mice were imaged for the complement strain per experiment. The mice were injected intraperitoneally with 5 mg of D-luciferin dissolved in 100 µL of PBS 10 minutes prior to imaging with an IVIS Spectrum live animal imaging system (Caliper Life Sciences, Hopkinton, MA), with the exception of one mouse that was infected with each *B. burgdorferi* strain tested but did not receive D-luciferin substrate. This mouse served as a negative control for background luminescence [60]. Imaging of the mice was performed 1 hour and at 1, 4, 7, 10 14 and 21 days post-infection [60]. After 21 days, the mice were sacrificed and the ear, abdominal skin, inguinal lymph node, heart, bladder, and tibiotarsal joint tissues were collected from each mouse aseptically for *in vitro* cultivation. Samples from ear, abdominal skin, inguinal lymph node, heart and tibiotarsal joint tissues were also collected from these mice for qPCR analysis of *B. burgdorferi* burden as described previously [60, 90].

### RNA Isolation for conventional RT-PCR and qRT-PCR

Three independent cultures of *B. burgdorferi* strains ML23 [59] (parent), the sRNA mutant strain DM103, and the genetic complement strain DM113, were grown to mid-log phase of 5 x10^7^ cells per ml at either mammalian-like conditions, defined as: 37°C, pH 6.8, under 5% CO_2_ or at conventional microaerophilic conditions of 32°C, pH 7.6, under 1% CO_2_. The cultures were centrifuged at 4500 x *g* for 20 minutes at 4°C, washed with 1 ml PBS, and centrifuged at 14,000 rpm for 15 minutes. The pellet was resuspended in 100 μl of sterile water and 300 μl of TRIzol (Invitrogen, Carlsbad, CA) was added prior to employing Direct-zol RNA Miniprep (Zymo Research, Irvine, Ca, USA) for total RNA isolation. The resulting RNA was treated with DNAse I (Roche, Indianapolis, IN) and RNAsin (Promega, San Luis Obispo, CA) to eliminate contaminating DNA and inhibit RNAse activity, respectively. To ensure that there was no contaminating genomic DNA in the cDNA reaction mixtures containing cDNA generated without reverse transcriptase were also included as controls. For conventional RT-PCR, 200 ng of total RNA from *B. burgdorferi* strains grown under conventional microaerophilic conditions were used to reverse transcribe into cDNA using primer sRNA R (Table S7) and SuperScript III (Thermo Fisher Scientific, Waltham, MA). Primers sRNA F and sRNA R (Table S7) were used to amplify *ittA.* For conventional RT-PCR of genes *bbd18* and *bbd21*, 1 μg of total RNA from *B. burgdorferi* strains grown under mammalian-like conditions were used for reverse transcription into cDNA using random primers (Thermo Fisher Scientific, Waltham, MA) and SuperScript III (Thermo Fisher Scientific, Waltham, MA). Oligonucleotide primers for *bbd18* and *bbd21* (Table S7) were used to amplify the genes.

For qRT-PCR analysis, 1 μg of total RNA from *B. burgdorferi* strains grown under mammalian-like conditions were used for reverse transcription into cDNA using random primers (Thermo Fisher Scientific, Waltham, MA) and SuperScript III (Thermo Fisher Scientific, Waltham, MA). Oligonucleotide primers from Table S7 were used to amplify specific *B. burgdorferi* strain B31 targets. Template cDNAs generated by the strains under the same conditions were normalized by using the constitutively expressed *flaB* gene as previously described [10, 18] with the ΔΔ*C_t_* method, in which the quantity of a given transcript is determined by the equation 2 ^- ΔΔ^*^Ct^*, where *C_t_* is the cycle number of the detection threshold.

### Northern blot Analysis

Northern blotting of SR0736/*ittA* was conducted using the same probe described previously [40, 102] from RNA extracted from *B. burgdorferi* strains grown under conditions that mimic mammalian-like conditions.

### DNA extraction of *B. burgdorferi* from infected tissues and qPCR analysis

Total DNA was isolated from ear, abdominal skin, inguinal lymph node, heart, and tibiotarsal joint using Roche High Pure PCR template preparation kit (Roche, Indianapolis, IN) as previously described [62,63,90]. Total DNA of 100 ng was used for each qPCR reaction. Quantitative real-time PCR analysis was conducted using the Applied Biosystems StepOnePlus Real-Time PCR system. *B. burgdorferi* genome copies and mammalian cell equivalents were determined using oligonucleotide primers in Table S7.

### RNA Sequencing (RNA-seq)

Three independent cultures of *B. burgdorferi* cells were grown to mid-log phase of 5 x 10^7^ cells per ml at 37°C, pH 6.8 and 5% CO_2_, e.g., *in vitro* conditions that mimic mammalian-like infection. Cells were centrifuged at 4,500 x *g* for 30 minutes at 4°C. Approximately 1 x 10^9^ cells were lysed in 1ml of TRIzol (Invitrogen, Carlsbad, CA) and RNA extracted following manufacturer’s instructions. RNA was checked for quality using the Agilent TapeStation 2200 standard RNA screen tape and quantified using the Qubit 2.0 Broad Range RNA assay. Total RNA was normalized between all samples for sequencing library preparation using the TruSeq Stranded Total RNA library preparation kit with ribosomal depletion. Each sample was uniquely barcoded, then the libraries were pooled at equal concentrations. Library pools were sequenced on a 2 x 75 bp paired-end sequencing run generating ∼15 million reads/sample by the Texas A&M Institute for Genome Sciences and Society (TIGSS).

After sequencing, a total of approximately 90.7 million 75 bp paired-end raw sequencing reads were checked to trim any adapter sequences and low quality bases using Trimmomatic [103]. Reads were scanned with a sliding window of 5 bp, cutting when the average quality per base drops below 20, then trimming reads at the beginning and end if base quality drops below 20, and finally dropping reads if the read length is less than 50. Thereafter, approximately 90 million filtered reads (99.2%) were mapped to the *Borrelia burgdorferi* strain B31 genome assembly (accession: GCF_000008685.2) using HISAT version 2.1.0 [104]. Average mapping rate was about 85.8% (Table S8). Transcript wise counts were generated using featureCounts tool from the SUBREAD package [105]. In total, the transcripts of 1,343 *B. burgdorferi* genes were tracked in this analysis. Differentially expressed genes were identified using a 5% False Discovery Rate threshold and a 2-fold change cut off with DESeq2 [106].

The data accumulated from the RNA-seq analysis are available at BioProject via accession number PRJNA565255.

### Tandem Mass Tags (TMT)

Three independent cultures of *B. burgdorferi* cells were grown to mid-log phase of 5 x 10^7^ cells per ml at 37°C, pH 6.8 and 5% CO_2_ (note: the cells used here were the same as those used for RNA-seq). Cells were centrifuged at 4,500 x *g* for 30 minutes at 4°C and washed twice with 10ml of cold PBS. Cells were resuspended in 50 mM triethylammonium bicarbonate (TEAB) and 5% SDS. Samples were quantified using Pierce BCA protein assay kit (Thermo Fisher Scientific, Waltham, MA) and 1.2 mg of protein was used for Tandem Mass tags (TMT) experiment. For this approach, protein extracts were isolated from the cells, reduced, alkylated, and proteolytically digested overnight. Samples were labeled with the TMT reagents in a 6-plex experiment and combined before sample fractionation and clean up. Labeled samples were analyzed by high-resolution Orbitrap LC-MS/MS. Identification and quantification of proteins was performed using Proteome Discoverer 2.2 software. The University of Texas Southwestern proteomics core performed the TMT analysis. The total of 718 *B. burgdorferi* proteins were detected in this analysis based on False Discovery Rates of less than 5%.

The TMT raw data was deposited to MassIVE using Proteomic Xchange Consortium with the data set number of PXD015685.

### SDS PAGE and immunoblotting

*Borrelia burgdorferi* protein lysates were resolved on a 12.5% polyacrylamide gel, transferred to a PVDF membrane, and blocked using non-fat powdered milk as done previously [90, 107]. Primary antibodies were used at the following dilutions: anti-Oms28 at 1:1000; anti-OspD at 1:5,000 and anti-FlaB at 1:5,000. Secondary antibodies with horseradish peroxidase (HRP) conjugates were used to detect immunocomplexes, specifically, anti-mouse Ig-HRP (Invitrogen, Carlsbad, CA, USA) or anti-rabbit Ig-HRP (GE Healthcare, Chicago, IL, USA) both diluted to 1:5,000. The membranes were washed extensively in PBS, 0.2% Tween-20, and developed using the Western Lighting Chemiluminescent Reagent plus system (Perkin Elmer, Waltham, MA, USA).

### Statistical analysis

For real-time qPCR analysis, multiple unpaired *t*-test, one per tissue, were performed to analyze the strains and corrected for multiple comparisons using the Holm-Sidak method. For quantitative reverse transcription PCR (qRT-PCR), one-way ANOVA was performed to analyze the strains. For the analysis of *in vivo* luminescence of mice, two-way ANOVA was performed. For the proteome volcano plot, multiple unpaired *t*-test, one for each protein identified in the 1% False Discovery Rate (637 proteins), were performed between the strains and corrected for multiple comparisons by using the Holm-Sidak method. Significance was accepted when the *p*-values were less than 0.05 for all statistical analyses employed.

### RNA target and structure prediction

The coding sequence of gene *bba66* was subjected to RNA binding predictions using IntaRNA (Freiburg RNA Tools; [93] against the processed *ittA* sequence. Mfold was used to predict the secondary structure of *ittA* [108].

### Ethics Statement

Animal experiments were performed in accordance to National Institute of Health (NIH) Guide for Care and Use of Laboratory Animals. Animal experiments also followed the guidelines of the Association for Assessment and Accreditation of Laboratory Animal Care (AAALAC). Approval for animal procedures was given by Tufts University and Texas A&M University Institutional Animal Care and Use Committees (IACUC; protocols B2018-98 [Tufts] and 2015-0367 [Texas A&M]). Mice were euthanized in manner that conforms to the guidelines put forth by the AVMA and was approved by Tufts University and Texas A&M University IACUC.

## ACKNOWLEDGEMENTS

We thank Lauren Weise, Parmida Tehranchi, Kristen Sanchez, and Alexandra Powell for excellent technical assistance for the work done at Texas A&M University. We would like to thank Andrew Hillhouse and Kranti Konganti from the Texas A&M Institute for Genome Sciences & Society (TIGSS) for their help with RNA-seq and the subsequent data analysis. We also want to extend our gratitude to the University of Texas Southwestern Proteomics core, specifically Andrew Lemoff and Mohammad Goodarzi, for their guidance through the TMT analysis.

## SUPPLEMENTAL TABLE DESCRIPTIONS

**Table S1.** Transcriptome of Parent and Mutant *ittA* strains. Deseq2 analysis was employed to the RNA-seq of Parent and the *ittA* mutant strains. A total of 1,343 transcripts were expressed in the Parent and the Mutant *ittA,* grown *in vitro* at mammalian-like conditions.

**Table S2.** Differentially expressed transcripts with a fold change of at least (+/- 1.4) in Parent versus the *ittA* mutant strain. Deseq2 analysis was employed to the RNA-seq of Parent and Mutant strains grown *in vitro* at mammalian-like conditions. A total of 92 transcripts were differentially expressed between the groups at the +/-1.4-fold change cutoff.

**Table S3.** Differentially expressed transcripts with fold change (+/- 2) in Parent vs versus the *ittA* mutant strain. Deseq2 analysis was employed to the RNA-seq of Parent and Mutant strains grown *in vitro* at mammalian-like conditions. A total of 19 transcripts were differentially expressed between the groups at the +/- 2-fold change cutoff.

**Table S4.** Differentially expressed transcripts with fold change in the range of (+/- 1.4 to 1.9) in Parent versus the *ittA* mutant strain. Deseq2 analysis was employed to the RNA-seq of Parent and Mutant strains grown *in vitro* at mammalian-like conditions. A total of 73 transcripts were differentially expressed between the groups at the +/- 1.4 to 1.9-fold change cutoff.

**Table S5.** Proteome of the Parent and Mutant *ittA* strains. A total of 718 proteins were identified by employing Tandem Mass Tags (TMT) proteomics technologies to Parent and Mutant strains, grown at mammalian-like conditions.

**Table S6.** Relative abundance of proteins with fold change of at least (+/- 1.4) in Parent versus the *ittA* mutant strain. A total of 69 proteins were identified with a fold change cutoff of +/- 1.4 and with an adjusted *p*-value of <0.05 between the groups.

**Table S7.** Oligonucleotides used in this study.

**Table S8.** Raw reads of the RNA-seq experiment between the Parent and the *ittA* Mutant groups. Three biological replicates of strains Parent and the *ittA* Mutant were grown *in vitro* at mammalian-like conditions and subjected to RNA-seq analysis.

## SUPPLEMENTAL FIGURE LEGENDS

**Figure S1.** The expression of genes *bbd18* and *bbd21* are not affected in the *ittA* sRNA mutant strain. The parent, sRNA mutant (*ittA*::Str^R^) and complement (Comp) strains were grown *in vitro* and total RNA was purified from each. Oligonucleotide primers specific for *bbd18* and *bbd21* were used without (-) and with (+) added reverse transcriptase (RT). The DNA ladder is shown at the left and the corresponding base pair values are indicated.

**Figure S2.** *In vitro* growth of the *B. burgdorferi* strains used in this study. The *B. burgdorferi* parent strain, the *ittA* sRNA mutant (*ittA*::Str^R^) and the sRNA complement strain (Comp) were grown in conventional microaerophilic conditions of 32°C, 1% CO_2_, pH 7.6 in triplicate in BSK-II media and enumerated by dark field microscopy daily out to day 9. No significant differences in growth were observed. Similar growth kinetics were observed between these three strains when the cells were grown at conditions of 37°C, 5% CO_2_ and pH 6.8 (data not shown). Data points shown reflect average value with standard error.

**Figure S3.** Overexposed Northern Blot reveals additional sRNA bands in the complement strain. Three biological replicates of the *B. burgdorferi* parent strain, the sRNA mutant (*ittA*::Str^R^) and the sRNA complement strain (Comp), were grown in mammalian-like conditions, RNA was purified and the *ittA* probe was used for Northern blot analysis at longer exposure. The *ittA* mutant does not expressed *ittA,* as expected. The stable processed form of *ittA* is observed as the dark band underneath the 100 nucleotide marker. The parent and complement strains expression of *ittA* is comparable, but between 800 and 300 nucleotides, the complement strain exhibits additional bands that are missing in the parent strain and could possibly contribute to the partial complementation of the strain *in vitro.* The marker is shown in nucleotides at the left of the blot.

